# Dissecting Genotype-Environment interactions with functional implications for parental selection in Cannabis Breeding

**DOI:** 10.1101/2025.04.17.649228

**Authors:** Anna Halpin-McCormick, Robert Thomson, Robert C. Clarke, Jeffrey Neyhart, Michael B Kantar

## Abstract

As climate variability continues to impact agricultural systems, identifying genetic factors that contribute to environmental adaptation will be essential for optimizing breeding strategies for the development of climate resilient varieties. Through human cultivation and naturalization, *Cannabis sativa* has dispersed globally, adapting to a range of environmental conditions across various climates and latitudes. We combined raw data from multiple public sources to conduct an Environmental Genomic Selection (EGS) analysis on 149 *Cannabis sativa* samples, to assess how different populations of Cannabis relate to their environmental conditions. Exploring Genomic Estimated Adaptive Values (GEAVs) across bioclimatic variables can facilitate the selection of parental material adapted for a specific condition. We further explore potential mechanisms of local adaptation by characterizing the individual marker effects which underlie these GEAV scores. To facilitate interpretation, we used previously described genetic groupings (Basal, Hemp-type, Drug-type feral, Drug-type). Distinct patterns emerged across population groups with the drug-type (Type I) group showing consistently narrow GEAV ranges, whereas the drug-type feral group showed a broader distribution, often having high GEAVs for precipitation variables. A key climate variable difference was seen in monthly average values, revealing a seasonal response to precipitation in drug-type feral samples. Exploring marker-effect differences between seasonal GEAVs indicated a response to seasonal precipitation in drug-type feral samples. As this samples are sourced from geographic regions that have seasonal monsoons, they may have traits conferring flood tolerance (water logging) that could be introgressed into other backgrounds. The basal group also exhibited broad GEAV ranges across several bioclimatic traits, indicating they may be a valuable genetic resource for introgression to enhance environmental resilience. These findings underscore the importance of incorporating diverse germplasm into breeding programs to improve Cannabis resilience to changing environmental conditions. EGS provides a fast method to enable climate-conscious parental selection while gaining mechanistic information. Ultimately, we hope that such a strategy could support the development of climate-resilient Cannabis varieties tailored to both current and future environmental challenges.

## Introduction

### History of Cannabis

Cannabis (*Cannabis sativa* L.) has been cultivated by humans for thousands of years and domesticated for many diverse purposes (food, fiber and medicine) (Clarke & Merlin, 2013) (Clarke & Merlin, 2016) (Hillig, 2005) (M. Ren et al., 2019) (G. Ren et al., 2021). Broadly Cannabis can be classified into two categories; non-drug varieties which contain less then 0.3% tetrahydrocannabinol (THC) and drug-type cultivars which have higher THC concentrations. While popularly known for its psychoactive effects, the rising demand in medicinal applications has significantly increased production (Klein, 2017). However, due to historical prohibition, Cannabis has not extensively benefited from the application of modern breeding methods (Barcaccia et al., 2020), despite its versatility as a multi-use crop (Schilling et al., 2020). With the end of prohibition in many regions worldwide, Cannabis is now undergoing a renaissance in research and innovation, driven by the application of modern scientific tools. Significant advancements have been made in genomics and the development of reference genomes (van Bakel et al., 2011) (S. Gao et al., 2020) (Braich et al., 2020) (McKernan et al., 2020) (Grassa et al., 2021), phenomics (Tran et al., 2024), transcriptomics (Guerriero et al., 2017) (Braich et al., 2019) and transformation techniques (X. Zhang et al., 2021). Despite these developments, much about this versatile plant remains to be discovered.

### Breeding History

Industrial hemp has become an established crop in many parts of the world, with a market valued at $258 million in the United States in 2023 (USDA National Hemp Report April 2024). Despite this, the history of breeding in Cannabis is poorly understood, with variable population structure seen across samplings and datasets (Sawler et al., 2015) (Lynch et al., 2016) (Soorni et al., 2017) (Henry et al., 2020) (G. Ren et al., 2021)(Halpin-McCormick et al., 2024) with evidence for natural and artificial selection (Woods et al., 2023) (G. Ren et al., 2021). Variation has also been observed in drug-type (Type I) cannabis varieties (Naim-Feil et al., 2021) (Lapierre et al., 2023) with genetic inconsistencies within strains of the same name also reported (Schwabe & McGlaughlin, 2019) and no clear genetic support for commonly referenced Sativa, Hybrid and Indica subcategories (Schwabe et al., 2021). Prohibition has limited the available information for Cannabis cultivars, regardless of population or group, leaving little understanding about the conservation status of wild or feral accessions and doubts about the existence of truly wild extant Cannabis (Small, 1975). This is further complicated by domesticated Cannabis escapees (feral Cannabis), which have been reported to revert to a wild-type phenotype within 50 years (Small, 1975) (McPartland & Small, 2020). Common hemp-type cultivars grown by farmers are usually maintained as Open-Pollinated Varieties (OPVs) and exhibit high within and between cultivar variability (M. Zhang et al., 2021) (Trubanová et al., 2023), with particularly large variation in flowering time (Stack et al., 2021). This can lead to performance variation for growers and subsequently market volatility. Understanding population structure in the genus can aid in the development of hybrid cultivars and breeding for uniformity, by reducing a large number of individuals into more manageable groups of relatedness or heterotic groups (Carlson et al., 2021).

### Species distribution and adaptation

A naturally dioecious wind pollinated plant, *Cannabis* has a wide variability in phenotypes which has complicated taxonomic assignments (McPartland & Small, 2020). A broad natural distribution, together with a long history of use by different human populations, has led to Cannabis becoming exposed and likely adapted to many different environments. It has been found in a large range of latitudes, altitudes and environments from temperate to monsoon, arid steppe and mediterranean climates (Clarke & Merlin, 2013) (Soorni et al., 2017) (Jin et al., 2020) (G. Ren et al., 2021) (Jehangir et al., 2024). Past ranges have also been examined converging on the northeastern Tibetan Plateau as a possible center of origin, with dispersions west and east (McPartland et al., 2019) (Rull, 2022), with multiple domestications likely (Long et al., 2017). However central Asian (Schultes, 1969) (Merlin, 1972) and East Asian (G. Ren et al., 2021) origins have also been proposed. Further, species distribution modeling suggests that there is likely to be broad loss of habitat suitability by 2050 due to changing climate (Halpin-McCormick et al., 2025). While populations exist that could thrive under these conditions, formal assessments examining the genetic relationship between the plant and the environment it inhabits in-situ are required (Babaei et al., 2024). Without substantial conservation efforts, these populations may experience adaptation lag, where adaptive evolution is unable to keep up with rapidly changing climates, resulting in population decline or local extinction (Wilczek et al., 2014).

There is extensive evidence demonstrating the influence of environmental factors on trait expression in crop systems (Chapin et al., 1993) (Harrigan et al., 2009) (Munkvold et al., 2013). In Cannabis, in experimental settings, adaptation to mountainous regions and altitude has been observed, resulting in significant changes in terpene and cannabinoid profiles (Pavlovic et al., 2019) (Cattaneo et al., 2021). Understanding how the environment affects plant secondary chemistry is particularly important for hemp producers to meet the regulatory requirements of maintaining below 0.3% THC at harvest. Cannabis plants exceeding this threshold of THC are classified as drug-type cannabis and subject to stricter legal controls. In addition, the growing medical and recreational markets require consistency in secondary chemistry. Geographical provenience of hemp should also be considered when selecting varieties with a specific end use to take advantage of local adaptation (Pavlovic et al., 2019). Hemp has promise as a drought-resistant fiber crop capable of thriving with relatively low water supplies (220 mm to 450 mm in a growing season) (Gill et al., 2023). A framework identifying functional traits which underlie the differential responses of crops to environmental variation can therefore help guide breeding programs and optimize crop attributes (Rolhauser et al., 2022). In order for expanding market demands to be met, it will be essential to breed climate-resilient Cannabis varieties that can thrive across diverse environments and latitudes.

### Identifying sources of local adaptation

As breeding for climate resilience gains importance (Cooper & Messina, 2023) many methods have been put forward to achieve this resilience across a range of species (Rellstab et al., 2015) (Ahrens et al., 2018) (Lei et al., 2019) (Todesco et al., 2020) (Waldvogel et al., 2020) (Neyhart et al., 2022). These efforts, have largely been related to GEA analysis (L. Gao et al., 2023), but this has yet to be done in Cannabis, with a broad species range and legal considerations presenting challenges to obtaining georeferenced samples.

A relatively new landscape genomic approach that can be used to select the best potential parents is Environmental Genomic Selection (EGS) (Campbell et al., 2024) (Halpin-McCormick et al., 2024). This technique makes use of environmental variables and genotype information to generate Genomic Estimated Adaptive Values (GEAVs) for climate potentialities. GEAVs represent the predicted genetic potential of an individual for adaptation to a specific environmental condition or variable. While identifying and collecting wild Cannabis samples can be complicated, the basal group identified in previous studies (e.G. Ren et al., 2021) could be considered a wild relative for the species. These, typically under-sampled, wild individuals have adapted to unique local climates and can provide valuable insights and useful alleles for developing more climate-resilient cultivars. Identifying local adaptation in cultivars is key for optimizing performance across diverse environments. Therefore, the objectives of this study were to extend EGS to combine both parental choice for breeding and the identification of potential mechanistic functions - in order to create a pathway to adapt *Cannabis* to current and future climates. To this end GEAVs were explored for seasonal and annual bioclimatic variables, with a specific focus on seasonal variation and the type of gene function enrichment that occurred when marker effects changed.

## Results

### Population Structure

Two data sets (Ren and Soorni) that contained whole genome sequence and geo-references were combined (G. Ren et al., 2021) (Soorni et al., 2017). We were able to recapitulate previously described population structure (**Table S1, S2, S5; Figure S1-S3**). Merging these two datasets yielded a total of 149 samples (**Figure S3**), 111 of which were georeferenced (**Figure 1A**). Principal component analysis (PCA) revealed group separation resulting in six distinct clusters which corresponded to previous phylogenetic assignments (**Figure S3C**). Genetic assignment using fastSTRUCTURE further supported these groupings with both the Elbow and Silhouette methods identifying k=5 as the optimal number of clusters (**Figure S4A**). Some of the samples falling in the basal grouping in PCA space displayed closer relationships to drug-type feral groups while others showed a stronger association with hemp-type samples (**Figure S4B**) and supported k=5. A rooted phylogenetic tree constructed from the shared SNPs between the two datasets (13,651 SNPs no missingness) showed similar group relationships (**Figure S5**) as seen in fastSTRUCTURE (**Figure S4**). We therefore categorized samples into Basal, Hemp-type, Drug-type feral, Drug-type, Population-1 and Population-2 groups in downstream analysis to better understand how these broad classifications may respond to varying environmental conditions.

**Figure 1.**
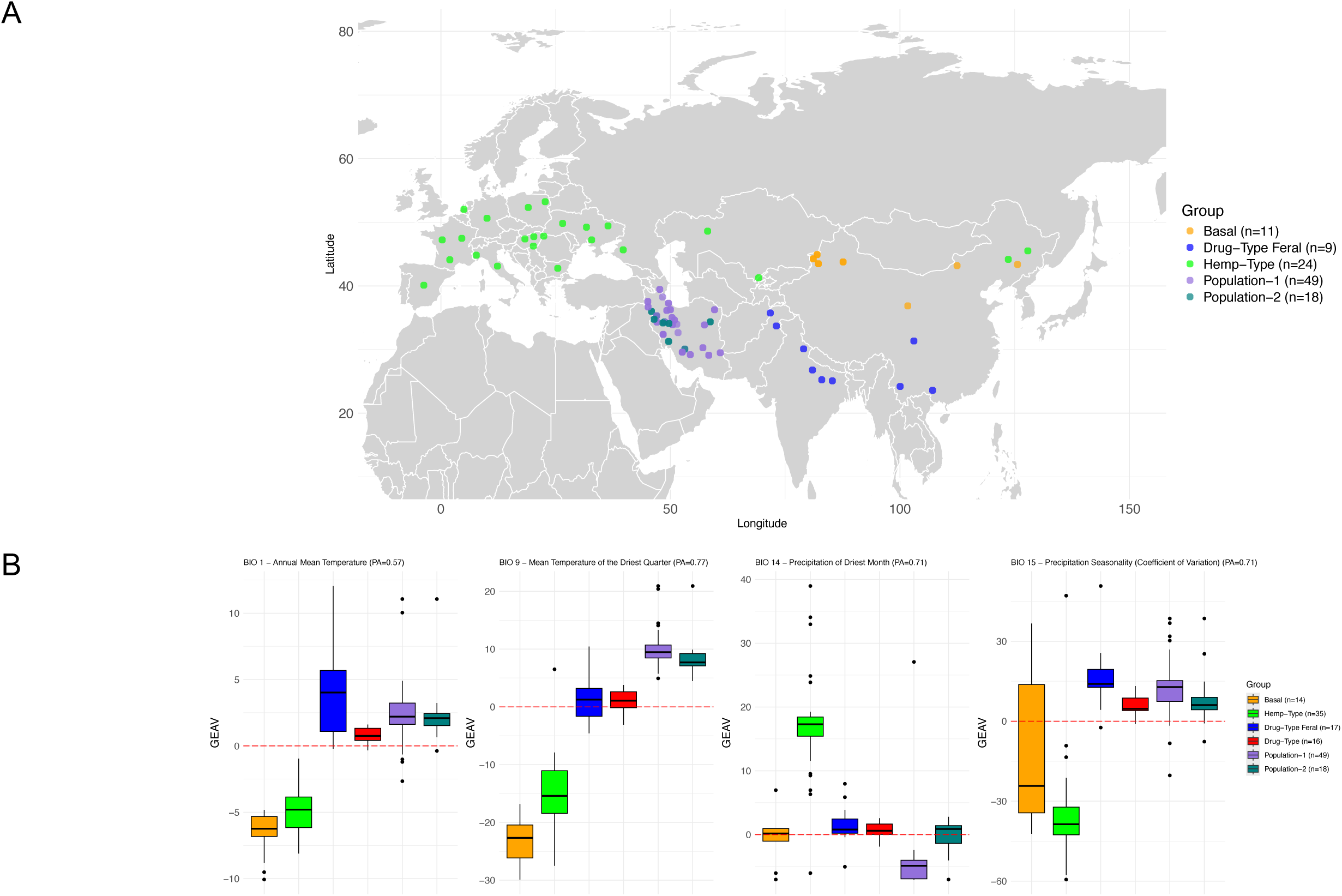
How group association relates to GEAVs for climate variables **(A)** Geographic distribution of the 111 georeferenced samples from Ren *et al* 2021 and Soorni *et al* 2017 **(B)** GEAV values for combined Ren *et al* 2021 and Soorni *et al* 2017 datasets (core n=50) estimated from 56,181 SNPs. The overall percentage of missing data was 4.2% and this missingness was imputed using the column mean for each SNP. Prediction accuracy (PA) for the rrBLUP model is indicated for each climate variable.

### Population climate variability and niche exclusivity

The raw climate experienced by each population revealed varying environmental conditions across groups (**Figure S6**). The distribution of the georeferenced samples revealed diversity across Köppen Geiger climate classes and population groups, with 15 of the 30 Köppen Geiger climate classes represented (**Figure 2**, **Figure 7A**). Some climate classes were observed commonly across all groups (e.g BSk - Arid desert, cold), while others were exclusive to one group (e.g. Dfb - Cold, no dry season and warm summer in hemp-type and Cwa - Temperate, dry winter, hot summer in drug-type feral). Thirteen of the fifteen environmental niches are represented in the core (**Figure S7B**) and ensures that training data encompasses a wide range of environments. A subset of environments contained more than five samples (**Figure S7C**) facilitating a more in-depth analysis. Redundancy Analysis (RDA) demonstrated that the combination of all 19 climate variables along with latitude, longitude and population structure explained 4.3% (adjusted R) of genetic variance (**Figure S14A, Table S8**). Further partitioning revealed that climate accounted for 1.48% of genetic variance after controlling for the effect of latitude, longitude and population structure (**Figure S14B**) while structure explained 0.03% of genetic variance when controlling for climate, latitude and longitude (**Figure S14C**).

**Figure 2.**
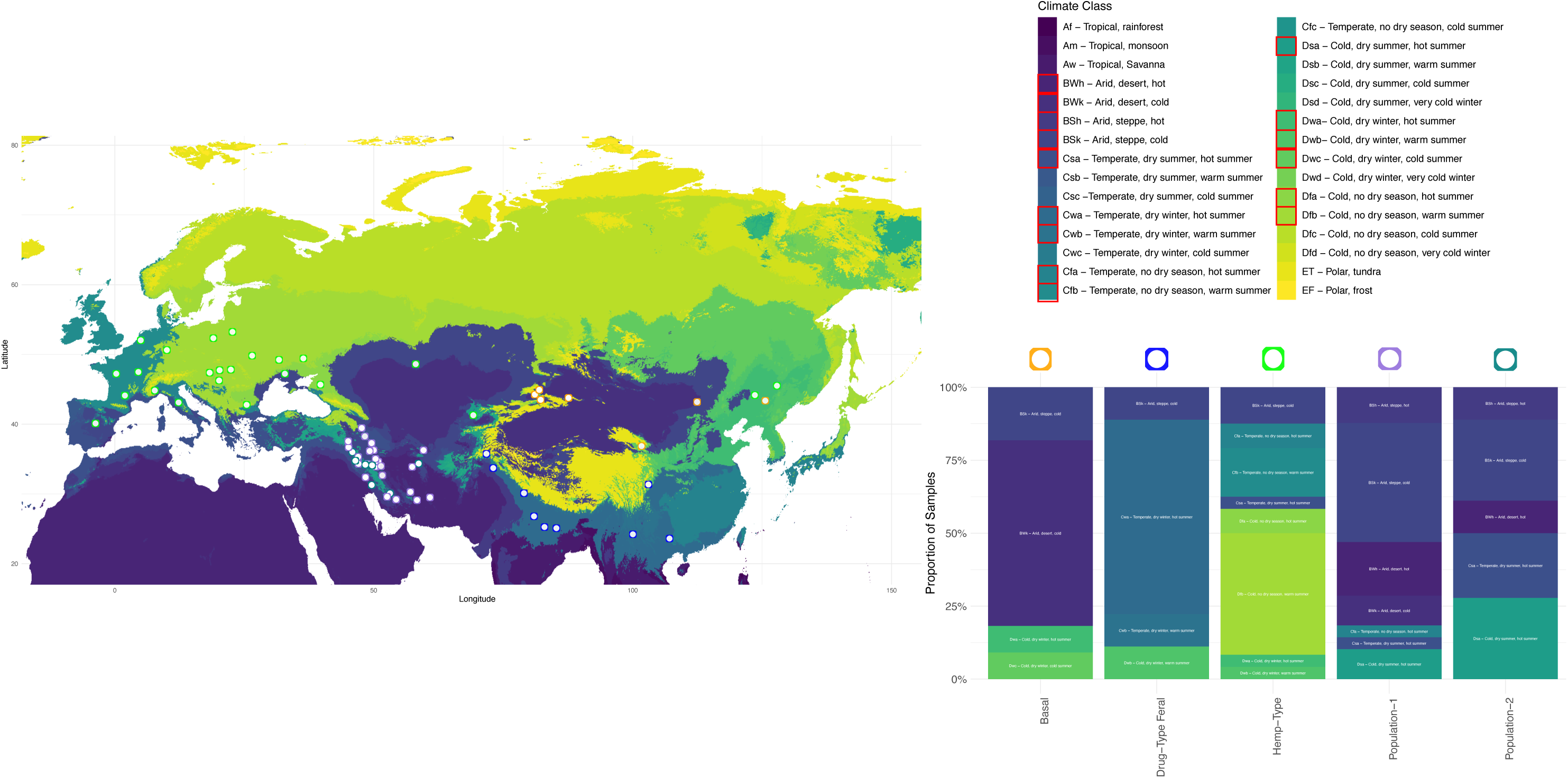
Geographic distribution of phylogenetic groups and Köppen-Geiger climate classes for the 111 georeferenced samples. Inset shows how the 15 climate classes covered are represented in each grouping.

### Genomic Estimated Adaptive Values (GEAVs)

To ensure representation of genetic diversity, a core set of 50 samples was developed using a greedy algorithm designed to maximize genetic distance across datasets (**Figure S3B, Table S3**). To minimize dataset bias we created a separate core of 25 samples from each dataset (Ren dataset - **Table S3** and Soorni dataset - **Table S6**) before combining them into the final core set (n=50) which was used as the training dataset (**Figure S3B, Table S6**). Genomic Estimated Adaptative Values (GEAVs) for each sample were calculated using climatic variables (Fick & Hijmans, 2017) providing insight into potential local adaptation. GEAVs are therefore analogous to predicted breeding values of an individual for adaptation to a specific environmental condition or variable. A similar trend in GEAVs and prediction accuracies was observed when comparing the results derived from the Ren dataset alone (∼ 2.9 million SNPs) (**Figure S8-S12, Table S4, S7**) to those obtained from the combined Ren and Soorni dataset (using 56,181 SNPs) (**Figure S10, S12, Table S7**). Population groups had distinct patterns associated with various climate variables **(Figure 1B, Figure S10, S12**). For BIO 1 (Annual Mean Temperature, Prediction Accuracy r = 0.57) we see high GEAV values for the drug-type feral, drug-type, population-1 and population-2 groups and these results are similarly observed for BIO 9 (Mean Temperature of the Driest Quarter, Prediction Accuracy r = 0.77). High breeding values for these traits indicate a potential tolerance to higher temperatures and during dry periods and contrast the negative values observed in the basal and hemp-type groups for these traits. For the precipitation-related variable BIO14 (Precipitation of the Driest Month, Prediction Accuracy r = 0.71) the hemp-type group displayed the highest GEAV, suggesting a potential adaptive advantage under minimal rainfall during the driest month. However, when examining BIO15 (Precipitation Seasonality, Prediction Accuracy r = 0.71) the hemp-type group exhibited a low GEAV mean which indicates limited breeding potential for seasonal precipitation. This is similarly observed when examining BIO 4 (Temperature Seasonality, Prediction Accuracy r = 0.73) where the basal group additionally has the highest GEAV mean (**Figure S10**). Across multiple climate variables the drug-type group consistently shows a narrow GEAV distribution (**Figure 1B, S10, S12**).

### Association between climate category and GEAVs

Seven climate classes contained more than five samples and facilitated further analysis (**Figure 3A**). Climate classes which fit this criterion included the following; Dsa (Cold, dry summer, hot summer), Dfb (Cold, no dry season, warm summer), Cwa (Temperate, dry winter, hot summer), BWk (Arid, desert, cold), BWh (Arid, desert, hot), BSk (Arid, steppe, cold) and BSh (Arid, steppe, hot). Within these, exclusive group assignments were chosen for downstream analysis as more clear relationships between group and climate could be examined (**Figure 3A**). By partitioning the GEAV values for individuals within these climate niches (**Figure S17**), individuals in the Cwa climate class showed consistently high GEAV values for precipitation variables (**Figure S17B**) and in particular for BIO12 (Annual Precipitation, Prediction accuracy r =0.56), BIO16 (Precipitation of Wettest Quarter, Prediction accuracy r =0.73) and BIO18 (Precipitation of Warmest Quarter, Prediction accuracy r =0.77) (**Figure 3B**). Genotype Environment Associations (GEA) were conducted for temperature related variables (BIO1-11) and precipitation variables (BIO12-19) using latent factor mixed models (LFMM) to identify possible large effect markers associated with environmental adaptation (**Figure S24 and S25**). Although a few markers were associated with climate variables, most of these were located in uncharacterized regions (**Table S9**).

**Figure 3.**
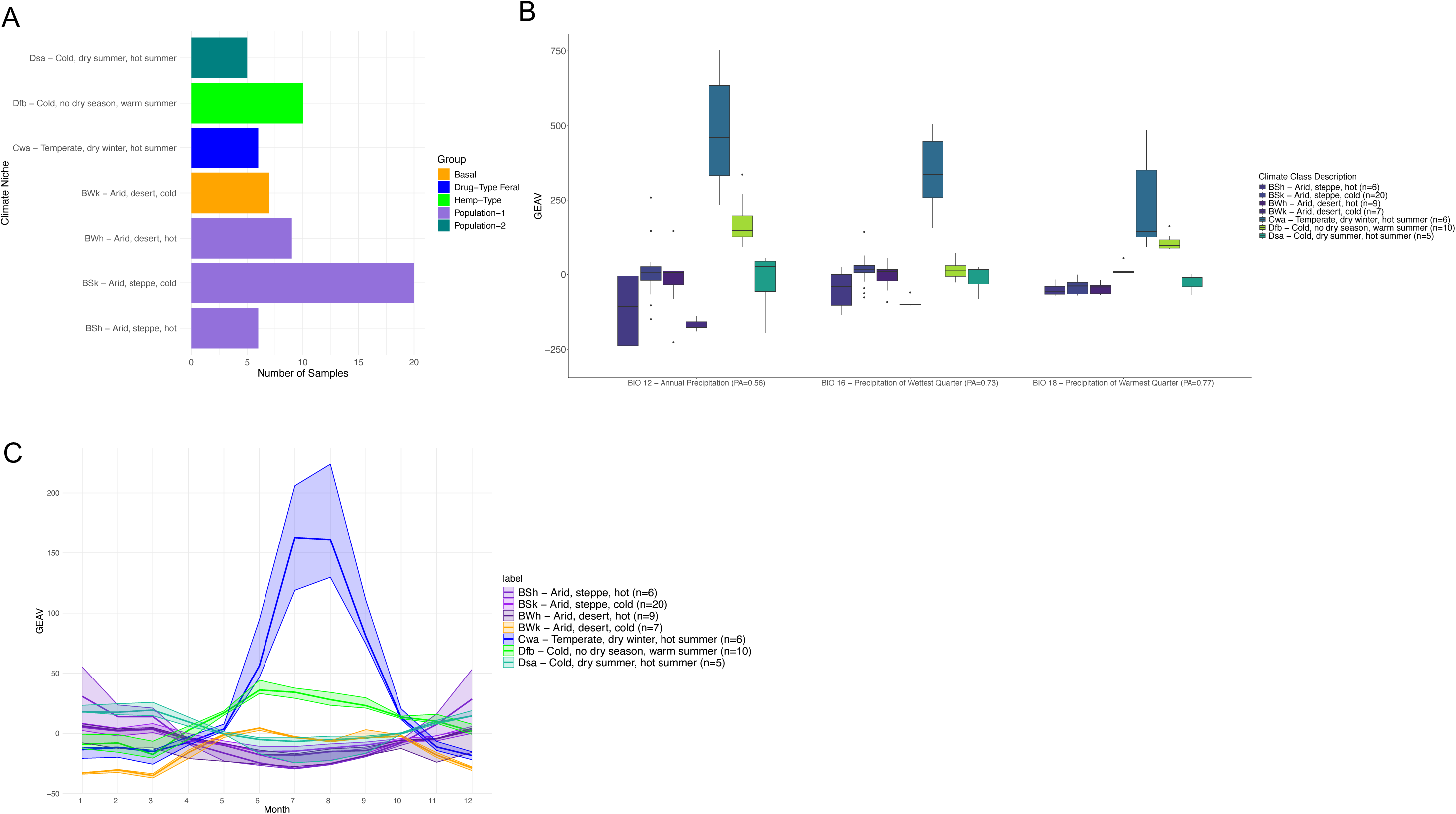
Relationship between environmental niches and GEAV values **(A)** Seven climate classes contained at least 5 samples from a single group **(B)** Consistently high GEAV values for the Cwa climate class (Temperature, dry winter, hot summer, n=6) for BIO12 (Annual Precipitation, PA=0.56), BIO16 (Precipitation of the Wettest Quarter, PA = 0.73) and BIO18 (Precipitation of Warmest Quarter, PA = 0.77) **(C)** Monthly averages (1970-2000) show seasonal response to precipitation in Cwa climate niche with GEAV values increasing from May to July/August.

### Seasonal Precipitation GEAV Response

Further exploring monthly averages for precipitation within these climate classes revealed a seasonal pattern for the samples within the Cwa climate class (n=6), most likely coinciding with the summer Monsoon season (**Figure 3C**). When examining seasonal responses over population groups, GEAVs were highest in summer in the drug-type feral group (n = 17) in both the Ren *et al* and the combined dataset **(Figure S19-22**). Seasonal fluctuation in GEAV values were observed, particularly for precipitation variables in the Cwa climate class, prompting further investigation into the markers which may be driving the marker effect size difference (**Figure 3C**, **Figure 4A**). We hypothesized that changes in GEAV means may be driven by a subset of markers whose effect sizes vary with environmental conditions throughout the year, akin to conditional neutrality, where some alleles may be favorable in one environment but neutral in another. Paired t-tests were conducted at each SNP site and across individuals to compare mean marker effects and determine significance between June and August **(Figure 4B).** A total of 788 SNPs showed a significant change in marker effect size (p = 0.01 after correction for multiple testing) **(Figure 4B**), with a large overall change in mean in marker effect size between June and August (p=2.2e-16) **(Figure 23C).** Of these 788 significant SNPs, 566 were located within genes in the cs10 assembly annotation (**Figure S23D**).

**Figure 4.**
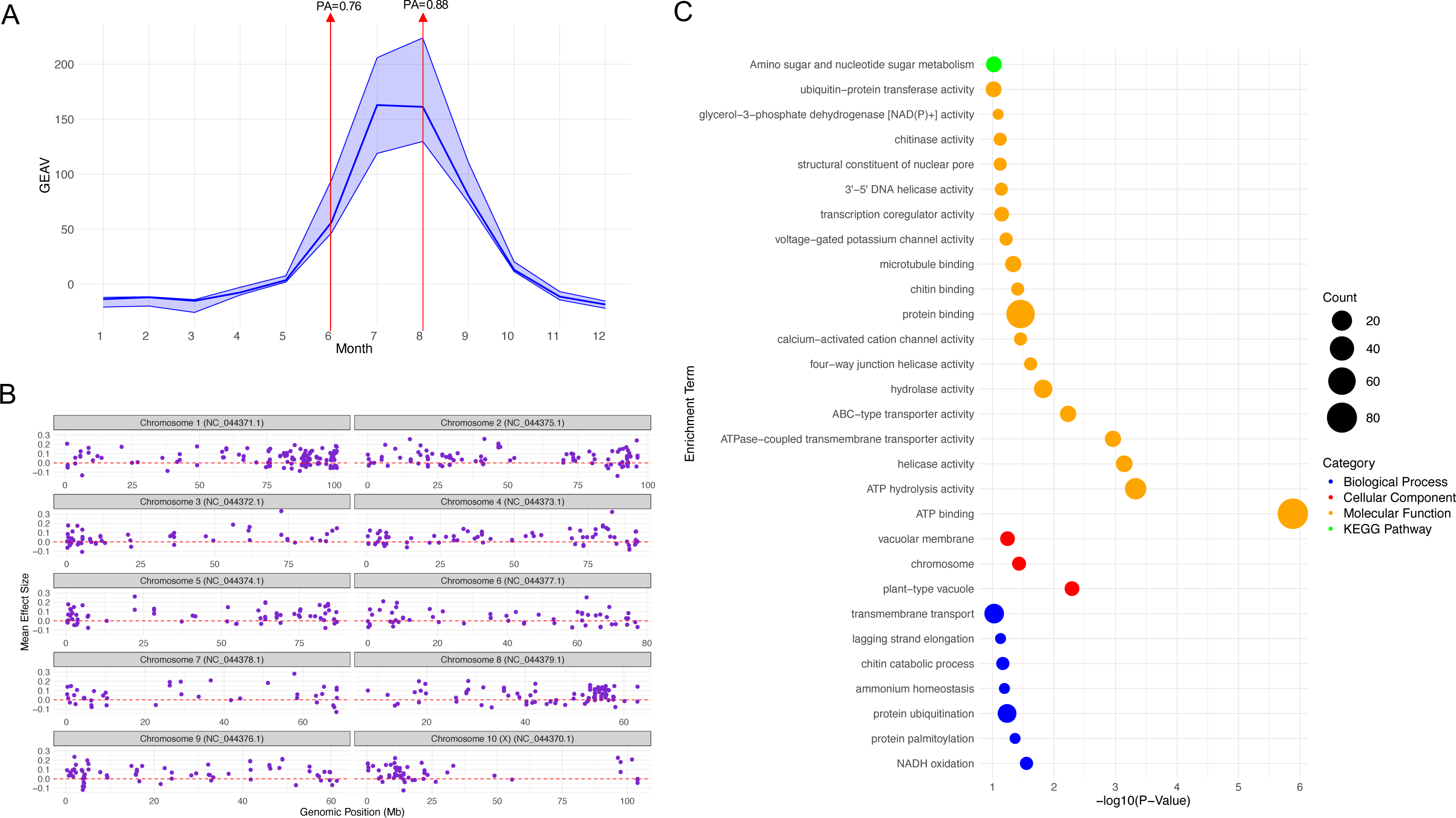
Analysis of seasonal response to precipitation in drug-type feral grouping **(A)** GEAV values for Drug-type feral (n=6) group show seasonal precipitation response which likely coincides with Monsoon season. The median line is plotted with the 25 and 75 interquartile range in shaded areas. Prediction accuracies were 0.76 and 0.88 for June and August respectively **(B)** Paired t-test at each SNP site across individuals between June and August for six individuals in the Cwa niche revealed 788 significant SNP sites which differed in mean effect size between timepoints. Mean effect size across individuals is plotted here for each of these sites for the month of August **(C)** The 788 significant SNPs represented 566 unique Entrez GeneIDs (ie. in some cases, multiple SNPs in the same gene region) and had 512 ortholog matches in *Nicotiana tabacum*. These 512 orthologs were then submitted for analysis using DAVID for enrichment in processes such as Molecular Function, Cellular compartment and Biological processes as well as KEGG pathway association. For reference a −log_10_(P-value) of 2 equates to a p-value of about 0.01.

Of these 566 genes, 512 has ortholog matches were found in *Nicotiana tabacum* using the OrthoDB software (https://gitlab.com/ezlab/orthodb_r/-/tree/main). Enrichment analysis using DAVID revealed a number of significant molecular functions at a −log_10_(P-value) of greater then 2 (p = 0.01) (**Table S10**). Enriched molecular functions included ATP binding (p = 0.0000013), ATP hydrolysis activity (p = 0.00047), Helicase activity (p = 0.00072), ATPase-coupled transmembrane transporter activity (p = 0.0011) and ABC-type transporter activity (p = 0.0059) (**Figure 4C)**. Additionally, an enrichment in plant-type vacuole (p = 0.0051) components is seen (**Figure 4C**), known to have roles in storage, detoxification and maintenance of cellular homeostasis through ion balancing and regulating of osmotic pressure within plant cells to control cellular turgor.

## Discussion

Our study successfully recapitulated population structure from previous work (G. Ren et al., 2021) (Soorni et al., 2017) (**Figure S1-S3**). Similar to previous work, we see the Iranian landrace types clustering closer to drug-type than to hemp-type (**Figure S3**; (Dehnavi et al., 2024)). Distinct group/population differentiation was evident with varying group GEAV means for bioclimatic variables (**Figure 1B, Figure S10, S12**). This highlights the importance of exploring the relationship between populations and their environments (**Figure 2**), as these interactions may drive advantageous allelic variation associated with specific environmental traits or properties. While we had cannabinoid content information available for some Iranian samples used here (Mostafaei Dehnavi et al., 2022) no associations with a distinct environment were observed (**Figure S7D**). This is not unexpected, as chemical profiles can be influenced by genetics, herbivory (Stack et al., 2023) (Park et al., 2019) as well as environmental properties (Yang et al., 2018) (Desaulniers Brousseau et al., 2021) (Pant et al., 2021) reported to have effects.

### Limitations

Genomic prediction models tend to perform better on larger datasets, with analysis on small sample sizes leading to poor prediction accuracy and poor effect size distributions. Here, we attempted to mitigate a small sample size by combining two datasets, which led to more individuals (n=149) but fewer genetic markers (56k). Additionally, a larger core (n=50) was developed to maximize genetic diversity while minimizing dataset bias. This was achieved by applying a greedy algorithm separately to the two datasets and then combining the results to capture as much of the diversity as possible across both datasets (**Figure S3B**). Imbalances in group sizes, such drug-type (n=14) versus hemp-type (n=35) samples, could additionally effect interpretations. Breeding history of a group is also important to consider, with hemp-type varieties often maintained as open pollinated populations, which may affect the strength of environmental associations. Prediction accuracies of each variable and timepoint should be considered (**Figure S8, 9, 18**) when considering GEAV values. There is a clear pattern between raw climate data (**Figure S6B**) and the group GEAV means (**Figure 1, Figure S12**). An additional limitation is the climate data for WorldClim is an average based on data from 1970-2000 and may lead to a less nuanced signal than would be desired. The annual life cycle of cannabis suggests that yearly averages may not effectively capture the strength of adaptive signals or more short-term environmental pressures. Consensus on Cannabis taxonomy remains a subject of debate. When applying “cultivated status” classifications, partitioning samples into feral, landrace and cultivated categories as reported in Ren et al., 2021 the differences in group separation (**Figure S1B**) and group GEAV means became less distinct (**Figure S11**). Classifying samples by cultivated status therefore introduces interpretative challenges. While the feral category displays more broad variability, it combines basal, drug-type feral and some hemp-type samples into a single group. Similarly, the landrace category merges basal and hemp-type samples. Grouping diverse populations into these three broad categories may obscure relationships between samples and their environments, as the inclusion of multiple climate classes and groups within each category reduces the clarity of environmental associations. Looking at markers that have changing effects due to season represents a novel way to try and explore mechanism while using a genomic prediction framework, however, these markers would benefit from experimental validation, potentially in controlled settings involving flooding conditions and transcriptomic analysis to confirm their presence and roles. Incomplete sampling of the genus is a concern in any analysis, including here. It is essential to carefully consider the limitations cited here when interpreting results.

### Detection of complex local adaptation using EGS and expanding to new Geographies

Genomic selection leverages the cumulative effects of numerous small-effect loci across the genome, with EGS extending the idea by combining genomic and environmental data to estimate the contribution of many small effect markers to an individual’s adaptive potential to a specific environmental trait. This approach is useful for identifying genetic adaptations to environmental conditions for polygenic traits. It provides different and complementary information from association analysis (**Figure S24-25**), or Fst outlier analysis (**Figure S26**), which focus on identifying individual large effect loci. Together, these strategies offer distinct advantages depending on the complexity of the trait being studied. Genomic variation across population groups and within specific environmental niches was examined (**Figure S15A-C**). These findings are consistent with previous studies which suggest that environmental adaptation is largely polygenic, with few markers of large effect likely to be found. Similarly, genome-wide Fst comparisons for each population within distinct climate classes against the basal group revealed some outlier markers (**Figure S26**). Examining nucleotide diversity across environmental groups reveals regions of lower diversity, in particular on chromosome 4, which may suggest a structural variation or inversion rather than an association with a particular niche (**Figure 15C**). Similar to the differences found here, previous work identified population differentiation on multiple chromosomes (1,3,4) (Woods et al., 2023) (Aina et al., 2025) suggesting chromosomal inversions as a reason for this (Lynch et al., 2024) (Dehnavi et al., 2024) (X. Chen et al., 2022). This may be an emerging genomic island of divergence as seen in other species (Renaut et al., 2013). Linkage disequilibrium (LD) analysis between hemp-type and population-2 in the 70-90 Mb region of chromosome 4 indicated that population-2 exhibits notably high LD across the 75-85 Mb region and higher then that seen in hemp-type. This pattern suggests the preservation of a non-recombining region or a more recent inversion in population-2 within this genomic interval compared to other populations (**Figure S16A/B**). Additionally, GEA analysis for BIO4 (Temperature Seasonality) (**Figure S24**) across all groupings shows a notable cluster of SNPs near the end of chromosome 4 with elevated -log10(p-values) and may suggest a genomic region of interest related to temperature seasonality. Individual SNP markers passed Bonferroni correction in this region of chromosome 4 additionally for BIO6 (Temperature of Coldest Month) and BIO11 (Mean Temperature of Coldest Quarter) (**Figure S24**) but corresponded to uncharacterized loci in the reference genome (**Table S9**).

The cultivation of crops necessitates a predictable timeframe from seed to harvest, making flowering time a critical trait of interest. Cannabis shows extensive flowering time variation (M. Zhang et al., 2021) (X. Chen et al., 2022). As climate changes, it might be necessary to move crops to different regions with suitable climates, exposing them to new photoperiods. This underscores the need to carefully select cultivars based on their suitability to the target growing environment, as local adaptation may render some varieties less compatible with different environmental conditions. The narrow distribution of the drug-type group may be due to limited sample size (n=16), but may also reflect a decoupling from broader environmental trends and may be related to the selective pressure this group has experienced, which has prioritized specific traits over environmental adaptability (**Figure S10, S12**). Drug-type varieties also represent a small subset of the genetic diversity within the genus which has been popularized and founder effects may therefore play a role in this narrow distribution and low GEAV ranges. Flowering time, or photoperiod sensitivity, is often governed by numerous genetic factors, with latitude often associated with variations in flowering time in a number of plant systems (Stinchcombe et al., 2004) (Debieu et al., 2013) (Xu et al., 2012) (González et al., 2021). Here examining GEAVs for latitude showed a strong correlation with realized latitudinal ranges (**Figure S13**). In this case, latitudes were unknown for the drug-type (n=16, Type I) samples in this dataset. Such an approach could aid in the selection of parent lines that are likely to align with the latitude (flowering time requirements) of the breeding program. Breeding for latitude-dependent photoperiod adaptation has greatest relevance for hemp production systems however, this trait is becoming increasingly important with growing interest in large scale outdoor cultivation of drug-type plants which have traditionally been cultivated indoors. In addition to flowering time, other agronomically important traits have received attention. For instance, traits like plant height, biomass, seed yield and biochemical QTL have been identified (Woods et al., 2021) as well as regions important for sex determination (Petit et al., 2020) (Shi et al., 2024). Incorporation of these large effect loci as fixed effects in prediction models may help improve model accuracies when examining GS of the trait, as has been seen in other crop systems (Spindel et al., 2016). Expanding the range will be dependent on flowering behavior and the tradeoffs between vegetative growth and reproductive phases (Carlson et al., 2021) (Dang et al., 2022). In addition to climate adaptation, there will also be demands to change flowering time to reach the necessary number of geographies where production is projected to be suitable. This means that breeding for flowering time, including auto flowering (day neutral) e.g. Autoflower 1, Early1 (Toth et al., 2022), Autoflower2 (Dowling et al., 2024), will also be important. To meet the growing demand and range of production environments, genomic approaches such as those proposed here enable an opportunity to readily select for multiple objectives simultaneously.

### Seasonal Genetic Pathway Enrichment

The significant enrichment for ATP binding/hydrolysis/helicase suggests a strong investment in ATP-related processes with roles in ATP binding, hydrolysis and ATPase-coupled transmembrane transport, as well as helicase and ABC-type membrane transporters which also use ATP in their functions. These processes in concert likely support stress adaptation mechanisms and aid in the seasonal response to precipitation, as energy-dependent membrane transport, DNA/RNA unwinding and the movement of molecules across cellular membrane, likely supports cellular detoxification and maintenance of ion gradients. Analysis of monthly averages from 1970-2000 revealed a seasonal response to precipitation in a subset of individuals namely; drug-type feral samples in the Cwa (Temperate, dry winter, hot summer) (**Figure 3C**). This variation in GEAV means across timepoints provided an opportunity to investigate the genetic markers contributing to this increase. Paired t-tests at each SNP site across individuals between June and August identified 788 SNPs with significant changes in mean marker effects (**Figure 4B**). These SNPs were mapped to 566 gene regions in the cs10 annotation and further analyzed for orthologs in *Nicotiana tabacum* to facilitate downstream pathway analysis (**Figure 4C**). Significant enrichment was observed for ATP-dependent activities, ATPase and ABC membrane transporters in molecular functions, while plant-type vacuoles showed enrichment in the cellular component category. We propose that this approach can provide valuable insights into the adaptive response exhibited by individuals for complex traits, in this example to seasonal precipitation. The changes in marker effect sizes in different seasons in specific environments coupled with an enrichment for genes with annotations related to stress adaptation in response to precipitation (**Figure 4**), suggests a prime mechanism for adaptation may be conditional neutrality (Lotterhos, 2023) (Tiffin & Ross-Ibarra, 2014). This relationship or conditional neutrality, becomes increasingly apparent in specific environmental contexts, such as conditions of increased precipitation and waterlogging seen here (**Figure 3C**, **Figure 4**). The detection of local adaptation to precipitation traits in a subset of individuals here, combined with the limitations of monthly and yearly averages, suggests that broader sampling and genotyping across diverse geographies, paired with local environmental data, could uncover stronger and more distinct signals of local adaptation within unique ecological niches. It is likely that many more climate classes containing Cannabis exist beyond those presented here (**Figure 2**), with recent species distribution modelling also supporting this (Halpin-McCormick et al., 2025). Recently, a comprehensive analysis using phylogeographic subgroups has revealed geographically defined groups within the genus (Balant et al., 2024). Further research is therefore needed to explore local adaptation within in-situ populations.

### Using EGS as a method for parental selection for exotic germplasm introgression

Climate change can be characterized by increases in temperature, drought and CO_2_ (Fawzy et al., 2020). Adaptation strategies to address these challenges include mechanisms around cell membrane stability, enhanced water content and improved photosynthetic capacity, as demonstrated in forages (Harmini & Fanindi, 2020). Improving breeding material through the use of exotic accessions is a well-known and often used procedure that can be accomplished using many different selection schemes (Iftekharuddaula et al., 2011) (Ru & Bernardo, 2020) (Wolfe et al., 2019). Exotic germplasm (e.g. landraces and wild relatives) often have increased stress tolerance and are commonly used in breeding (Hancock et al., 2011) (Prasanna et al., 2021) (Quezada-Martinez et al., 2021) (Nyoni et al., 2023). However, adaptational lag remains a critical concern in conservation and management decisions for many species (Wilczek et al., 2014). Exploring climate adaptation during the breeding process provides ample opportunity to identify rare recombination events that may have more beneficial phenotypes (Agha et al., 2024).

Environmental genomic selection offers an approach to identify parental lines from diverse geographic regions, which may aid in trait improvement in response to projected climate change. By ranking individuals by their GEAVs, for example tolerance to extreme temperatures (BIO5 or BIO10) direct selections could be made for parents for drought conditions or water use efficiency. The distribution ranges seen for the basal and drug-type feral groups over a variety of bioclimatic traits suggests that there is great potential for their use in breeding. Low seasonality GEAVs for hemp-type may reflect the groups tendency to inhabit temperate regions with relatively stable, no dry season, warm summer niches (**Figure 2**). The drug-type feral group however had the highest GEAV mean for BIO15 and suggests a possible adaptive relationship with seasonal precipitation. Interestingly, the basal group exhibited a broad GEAV distribution for BIO15, despite a low overall mean. This suggests higher variability within this group (**Figure 1B**) and may be related to their shared admixture with other classes as seen in fastSTRUCTURE analysis (**Figure S4**). Additionally, the approach taken here could be integrated with speed breeding protocols (Schilling et al., 2023) to decrease lag time in selection schemes for complex traits. There is currently significant untapped potential for crop improvement in Cannabis for a variety of uses, with the deployment of genomic tools for rapid selection still largely underutilized (Smart et al., 2022). Recent surveys have highlighted challenges in the hemp industry, with stakeholders desiring the production of stable and uniform cultivars with regional adaptability (Ellison, 2021). Work in other crop systems, suggests that breeding for local and broad performance are mutually supporting goals, with the development of broadly excellent and locally exceptional varieties aligning with breeder and grower needs (Ewing et al., 2024). Additional critical areas for research include improved pest resistance and disease management, with successes more recently in the identification of large effect loci important for powdery mildew (PM) resistance (Stack et al., 2021) (Mihalyov & Garfinkel, 2021) and susceptibility (Pépin et al., 2021) (Stack et al., 2024) (Seifi et al., 2024). Additionally, 82 other genes have also been associated with powdery mildew resistance (McKernan et al., 2020), indicating that PM manifestation is influenced by both large and cumulative marker effects.

## Conclusion

Cannabis representatives were seen across 15 distinct climate classes, with the possibility of more existing but yet to be sampled. Although some groups exhibited limited genetic variation with respect to climate, the variability seen within the genus suggests significant potential for adapting Cannabis to future climate challenges. The broad geographic distribution and GEAV values observed in basal and drug-type feral groups across climate classes highlights their utility as exotic germplasm in breeding programs. These groups have potential for introgression of climate resilient genes into other varieties and are therefore in need of conservation as well as ethical and equitable management. Drug-type feral samples demonstrated a seasonal response to precipitation, suggesting that local adaptation signals should be considered when selecting material, as this may challenge trait expressions in new environments. Resilience in cannabis could be envisioned as a spectrum across populations and environments, influencing its adaptive capacity to climate change. In order to combat the concerns of the environmental footprint of indoor cultivation (Summers et al., 2021) (Ashworth & Vizuete, 2017), attempts to move towards outdoor production by breeding for climate resilient varieties will be required and could reduce costs substantially. An aim that could be facilitated with methods such as those proposed here.

## Materials and Methods

### Data acquisition and processing

Raw reads for 82 samples were downloaded from NCBI under the project ID PRJNA734114 (G. Ren et al., 2021) using ENA ftp downloader (https://github.com/enasequence/ena-ftp-downloader) (**Table 1**). As this data was previously published, we as closely as possible tried to perform the same pipeline to maintain consistency. Adapter sequences were removed using fastp (S. Chen et al., 2018) (script_1_fastp). We used the CBDRx (cs10 v.2.0) (GCF_900626175.2_cs10_genomic.fna) as the reference genome (Grassa et al., 2021) (script_2_bwa_index). Reads w(Li & Durbin, 2010)a (v0.7.18) (Li & Durbin, 2010). The resulting SAM file was converted to BAM format using samtools (v1.12) (Danecek et al., 2021) with a quality threshold of 10 (script_3_map). Duplicate reads were removed from the BAM file using MarkDuplicates in Picard (http://broadinstitute.github.io/picard) (script_4_dedup). Labeling of read groups was corrected using AddOrReplaceReadGroups in Picard (v3.1.0) (script_5_add_RG). The reference genome was indexed using samtools faidx and a sequence dictionary was created for integration with GATK (DePristo et al., 2011) (script_6_gatk). Using RealignerTargetCreator and IndelRealigner in Genome Analysis Toolkit (GATK) (v3.8) (DePristo et al., 2011) local realignment around insertions and/or deletions (indels) was performed(script_7_realign_indel). Using bcftools (v1.11) (Danecek et al., 2021) BAM files were merged and variant calling was performed (script_8_mpileup). For filtering, variants were filtered using bcftools filter with a mapping quality of 30 (MQ < 30). Additional filtering steps were performed using with VCFtools (v0.1.16) (Danecek et al., 2011) and included the following criteria (i) restriction to biallelic variants (--max-alleles 2), (ii) setting a minimum read depth (DP) between 4 and 50 (--minDP 4, --maxDP 50), (iii) filtering out genotypes with a genotype quality (GQ) below 10 (--minGQ 10), (iv) excluding variants with a quality score (QUAL) below 30 (--minQ 30), (v) removing variants with a minor allele frequency (MAF) below 0.05 (--maf 0.05) and (vi) retaining variants only if present in at least 70% of samples (--max-missing 0.7) (script_10_filter). This resulted in in a total of 15,415,881 SNPs (pre-filtering was 51,570,415). Use type for each sample was assigned based on that previously published in Supplemental Table 1 (G. Ren et al., 2021) where “cultivation status” and “phylogenetic group association” were listed for each sample (here **Table S1**). Raw reads for 67 samples were downloaded from NCBI project ID PRJNA419020 (Soorni et al., 2017) (**Table 5**) and went through the vcf pipeline above. This resulted in in a total of 70,848 SNPs (pre-filtering was 1,209,397). Previous analysis of these 67 samples by authors showed clear population structure into two distinct groups (Soorni et al., 2017). With the exception of two samples, this population structure of two distinct groups was also observed here (**Figure S2, Figure S3).**

### Intersection of Datasets

These two datasets where then intersected for shared SNPs using bcftools isec resulting in 56,181 common variants, which post LD=0.2 resulted in 9,759 SNPs and population separation (**Figure 1**).

### Core development

For the Ren *et al* dataset, a core (n=25) was developed using all 15,415,881 SNPs. Using the scikit-allel (Pedregosa et al., 2011), NumPy (Harris et al., 2020) and Pandas (McKinney, 2010) python libraries, genotype matrices and genetic distance matrices were calculated. Georeferences were available for 44/82 samples (**Table S2**) and core development was therefore restricted to the samples with georeferences. Genetic distances were calculated using the Hamming distance metric though the pairwise_distances function in the scikit-learn library. To ensure maximum genetic distance between samples a greedy algorithm was applied (**Table S3**) (1_step1_core.py). A core of 25 was also developed for the Soorni *et al* dataset using all 70,848 SNPs. As all 67 samples in this dataset had georeferences, all samples were included for core development. Genetic distances were similarly calculated using the Hamming distance metric though the pairwise_distances function in the scikit-learn library, with maximum genetic distance between samples ensured using a greedy algorithm (1_step1_core.py). Downstream genomic selection on the fusion of these datasets used a core of 50 (**Table S6**) summed from both of these independent runs for each dataset. This was to prevent dataset biases and representation within the core.

### Environmental Data

Occurrence points were used to query the WorldClim 2.1 climate data. The standard 19 bioclimatic variables for temperature and precipitation (average for the years 1970-2000) and monthly averages (for years 1970-2000) for average temperature (tavg °C), minimum temperature (tmin °C), maximum temperature (tmax °C), precipitation (mm), water vapor pressure (vapr kPa), wind speed (wind m/s), solar radiation (srad kJ/m^2^/day) and elevation (m) were acquired. Data was downloaded at the highest available spatial resolution of 30 seconds (∼1 km^2^) (https://www.worldclim.org) (Fick & Hijmans, 2017).

### Environmental Genomic Selection

For the Ren *et al* dataset, prediction accuracy and genomic selection were performed using 2,896,658 SNPs as this subset of SNPs were found in 100% of samples. Genomic estimated adaptive value (GEAV) for each accession for a given trait (conceptualized as the genetic value for a specific environmental context (Cortés et al., 2022) were calculated (**Table S4**). Four genomic prediction methods were examined: (i) RR-BLUP, (ii) G-BLUP with an exponential kernel, (iii) G-BLUP with a Gaussian kernel, and (iv) BayesCπ. R packages ‘rrBLUP’ (Endelman, 2011) and ‘hibayes’ (Yin, L., Zhang, H., Li, X., Zhao, S., & Liu, 2022) R were used for the analysis (**Figure S8**). The training population (core n=25) consisted of 25 georeferenced accessions. The remaining **57** accessions were used as the validation set. Prediction accuracy was based on Pearson correlation (r(PGE,y)) between the predicted genotypic effects and the observed environmental variable with 10-fold cross-validation. Prediction accuracies and genomic selection was performed on the combined Ren *et al* and Soorni *et al* datasets similar to above using the fusion core (n=50) (**Table S6**) and using the 56,181 shared common SNPs. This core served as the training population with the remining 61 samples which had georeferences and therefore climate data used as the validation set. GEAVs for each accession and trait were calculated (**Table S7**). Due to ∼4% missingness across SNPs imputation using the mean at missing sites was required for genomic selection.

### Köppen-Geiger climate classes

Present day Köppen-Geiger climate classes were acquired at 1km resolution (Beck et al., 2018). Climate classifications for georeferenced samples were determined by intersecting each samples latitude and longitude coordinated with the Köppen-Geiger climate dataset.

### Phylogenetic and fastSTRUCTURE analysis

To examine structure and shared admixture proportions, VCFtools was used to generate map and ped files for the input VCF file. PLINK (v1.90b6.21) (Purcell et al., 2007) then converted these files into bed, bim and fam files. Population structure was then assessed using fastSTRUCTURE (v1.0) (Raj et al., 2014). The optimal K was examined using the silhouette and elbow methods in the FactoMineR (v2.11) (Lê et al., 2008) (Kassambara & Mundt, 2016)0.7) (Kassambara & Mundt, 2016) packages.

A maximum likelihood phylogeny was constructed on the 149 samples rooted using the *Humulus lupulus* reference genome (GCA_963169125.1_drHumLupu1.1_genomic.fna) (Natsume et al., 2015). The merged_Ren_Soorni_snps.vcf.gz was filtered for no missingness resulting in 13,669 SNPs. These SNP positions were mapped to corresponding positions in the *Humulus lupulus* genome using bedtools (v2.31.1) (Quinlan & Hall, 2010) and the corresponding SNPs extracted. To convert the vcf file to phylip format the vcf2phylip software was used (Ortiz, 2019) handling heterozygous ambiguities using the -m 1 mode and IUPAC ambiguity codes. A phylogenetic tree was inferred using IQ-TREE (v2.3.6) (Nguyen et al., 2015) with the best substitution model selected using -m MFP flag (TVM+F+ASC+R5) with ascertainment bias +ASC and -bb flag for bootstrap support. Trees were visualized in FigTree (v1.4.4) (http://tree.bio.ed.ac.uk/software/figtree/).

### Environmental contribution to dataset genetic variance

RDA analysis was performed in R studio using vcfR (v1.15.0) (Knaus & Grünwald, 2017) and adegenet (v2.1.10) (Jombart & Ahmed, 2011) to create a genotype matrix file from the VCF file (merged_Ren_Soorni_snps.vcf). Missing genotype data (∼4%) were imputed with the most common value for each SNP. The vegan (v2.6.8) (Oksanen et al., 2001) and psych (v2.4.6.26) (Revelle, 2007) packages were used to perform RDA, analyzing the contributions of all climate variables, latitude, longitude and population structure on genetic variance. Additional RDA analysis was conducted on partitioned subsets of climate variables, latitude, longitude and population structure to assess variance contributions (**Table S8, Figure S14**).

### Genome wide scans

Nucleotide diversity (pi) was calculated for each population to assess genetic variability. VCF files were subset by population using bcftools view with population specific sample lists for the BSh, Bwh, Bwk, Bsk, Dsa, Cwa and Dfb climate niches. Nucleotide diversity was then calculated at each site using vcftools --site-pi. Output was visualized using ggplot2 (**Figure S15**).

Latent factor mixed model (LFMM) analysis for Genotype Environment Associations (GEA) was performed to identify possible large effect SNPs associated with environmental variables. This was performed using all 56,181 SNP sites in all 149 samples. Using VCFtools the genotype data was extracted from the vcf file and converted to a numeric matrix with missing values imputed with the mean. GEA was performed using the lfmm package (v1.1) (Jumentier, 2020) in R studio using the lfmm_ridge function with k=5 to account for population structure. Individual SNP p-values were calculated and a Bonferroni threshold and false discovery rate (FDR) adjustment was applied to identify significant SNPs (**Figure S24/25**).

Genetic differentiation between populations using Fst analysis with the Weir and Cockerhams method. For each pairwise comparison, VCFtools was used to calculate Fst values between population pairs. In this case the basal group (in the Bwk niche) was compared to the Hemp group (in Dfb), Population 1 group (in BSk, Bsh and Bwh) and population 2 (in Dsa). By comparing each population to the basal group, the degree of genetic divergence in the other populations could be assessed relative to this foundational group. This approach helps identify specific genomic regions contributing to adaptive differences among populations across distinct environmental niches (**Figure S26**).

### Seasonal SNP effect size analysis and Ortholog Identification

A genotype matrix in numeric format was generated using vcfR only retaining SNPs without any missing data (13,670 SNPs). Genomic prediction accuracy and genomic selection were performed similarly to that described above. In the case of the seasonal trend observed for precipitation, samples were assessed for per SNP and individual marker effect sizes for the months of January and August. The per SNP and sample results were then partitioned to the Cwa climate niche (n=6) as this group appeared to have high GEAV means when examined across monthly averages. Significant SNPs with effect size differences between January and August were identified via paired t-tests with Benjamini-Hochberg (BH) correction (p>0.01).

To evaluate the impact of population structure, we compared marker effects estimated with and without the inclusion of principal components. Pearson correlations were high (r = 0.964 for August, r = 0.936 for June comparisons) suggesting minimal bias due to structure. Therefore, we proceeded without additional correction. SNPs with significant effect sizes between the two timepoints were annotated by overlapping SNP-IDs with genomic features from the cs10 GFF file. Unique gene identifiers were queried using OrthoDB in R Studio (v4.4.0) for *Nicotiana tabacum* orthologs. This list of orthologs was then submitted to DAVID (Sherman et al., 2022) selecting to identify potential functional enrichment for Molecular Function (MF), Cellular Component (CC), Biological Process (BP) and KEGG pathways.

## Data availability

Data and scripts are available at https://github.com/ahmccormick/Cannabis_EGS and at Figshare https://figshare.com/authors/Anna_H_McCormick/17741367.

## Author Contributions

**Conceptualization**: AHMC and MBK, **Formal Analysis:** AHMC, **Figure Preparation:** AHMC, **Manuscript writing and Revising**: AHMC, RT, RCC, JN, MBK.

## Acknowledgements

We would like to thank the University of Hawaii High Performance Compute Cluster.

## Supplemental Figures

**Figure S1.**
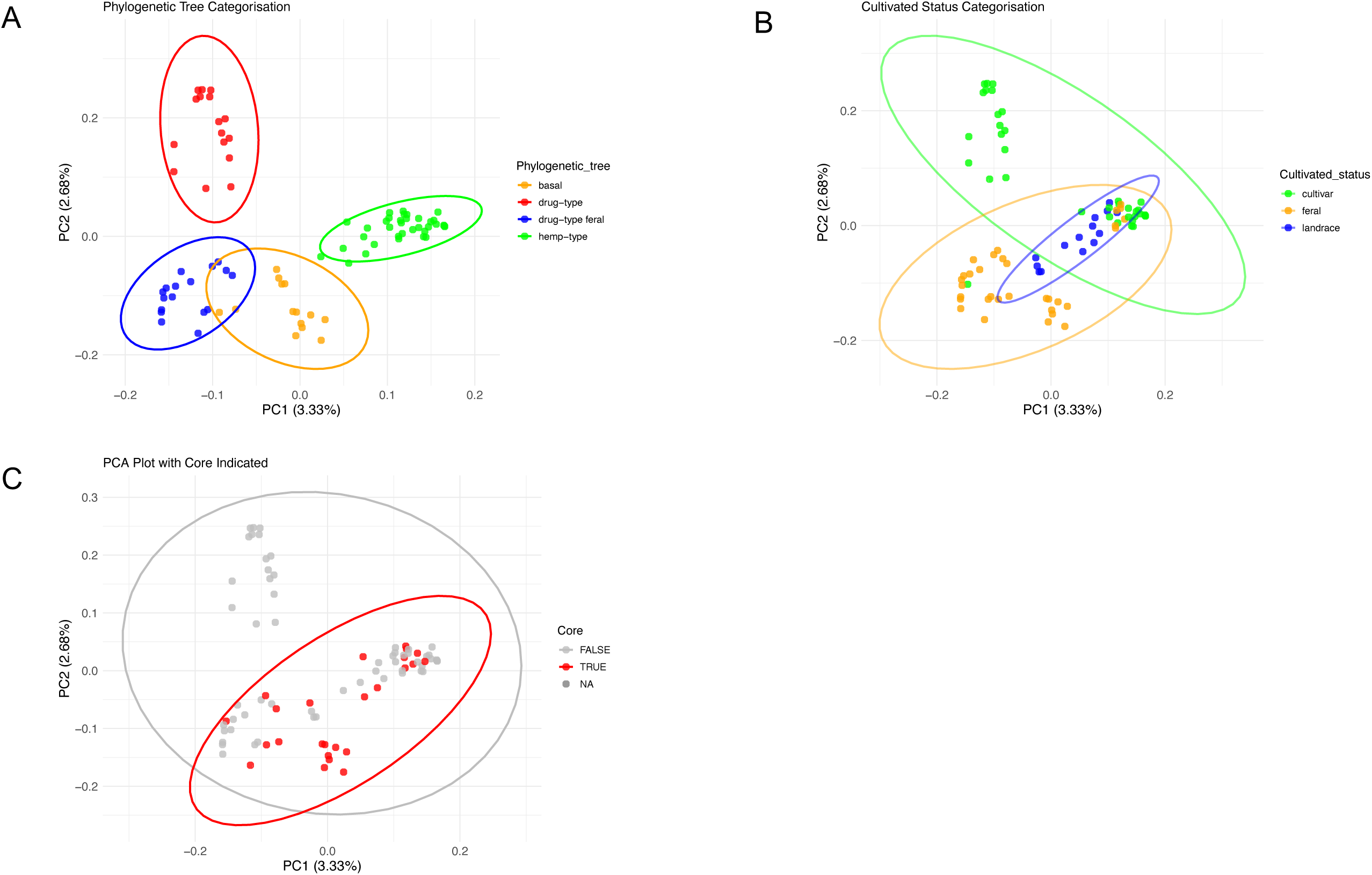
Recapitulation of the groupings previously published by Ren *et al*., 2021 **(A)** Distribution of lines in Principal Component space from 56,920 SNPs (LD=0.2) indicating phylogenetic associations **(B)** Cultivated status association and (C) Core affiliation (n=25) within the dataset.

**Figure S2.**
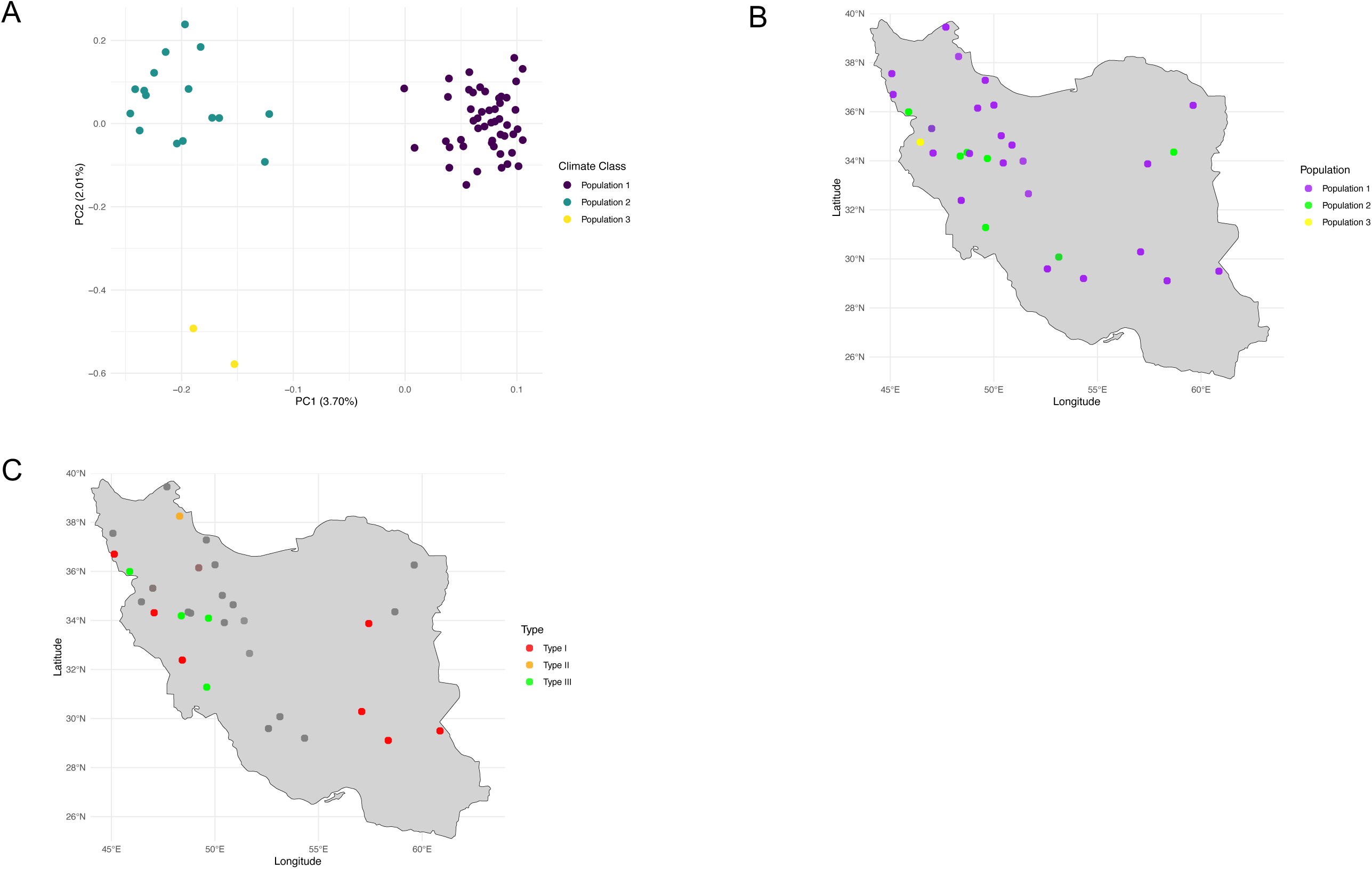
Groupings observed in the Soorni *et al.,* 2017 dataset **(A)** Distribution of lines in Principal Component space from 9,568 SNPs (LD=0.2**) (B)** Relationship between population and geographic location **(C)** relationship between secondary chemistry type and geographic location.

**Figure S3.**
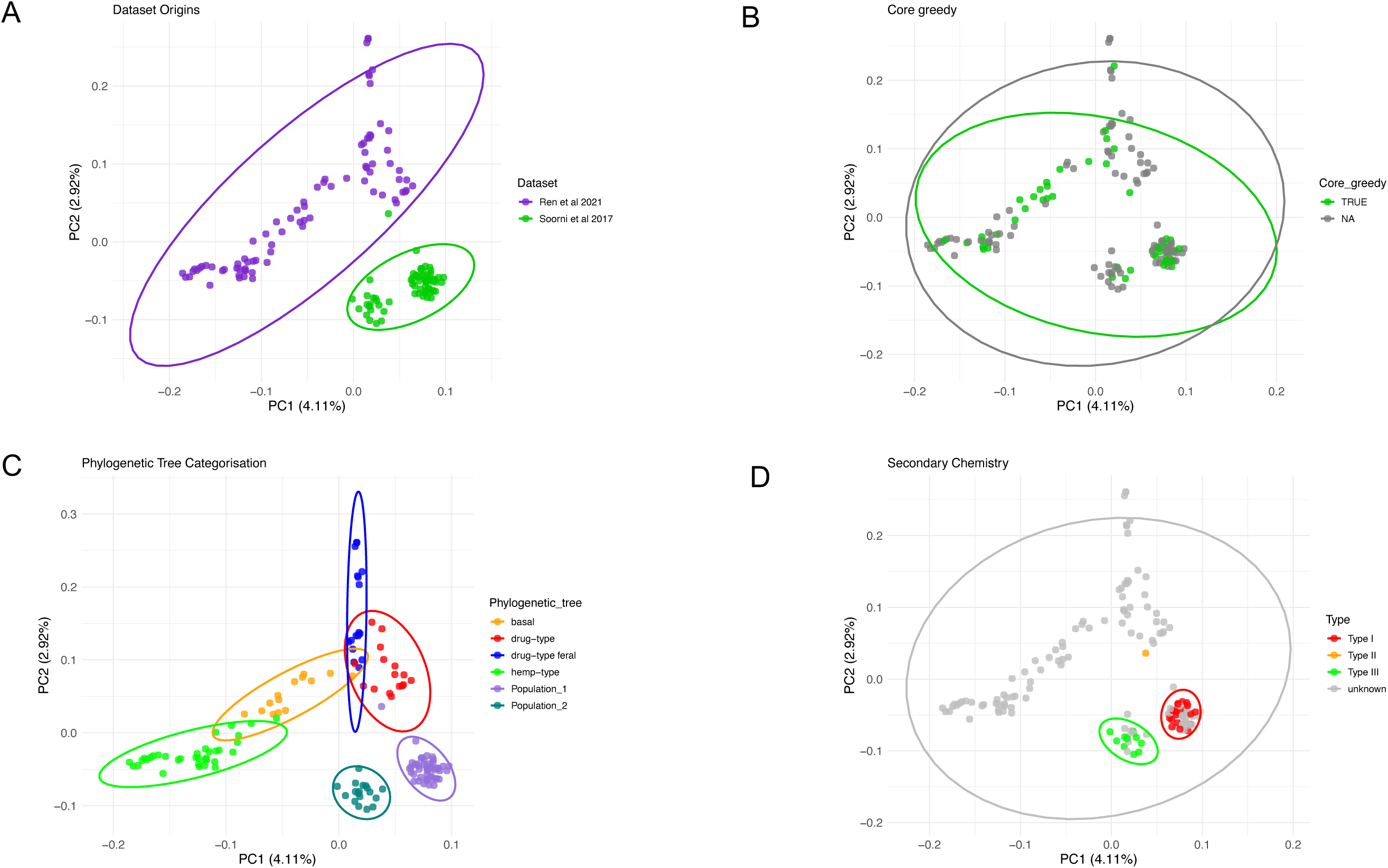
Principal component analysis (PCA) showing distribution of the 149 lines for the combined dataset based on 9,759 SNPs (LD=0.2) for **(A)** Dataset origin **(B)** Core affiliation (n=50) and **(C)** Phylogenetic associations effect distribution ranges **(D)** Distribution of samples with known chemistry.

**Figure S4.**
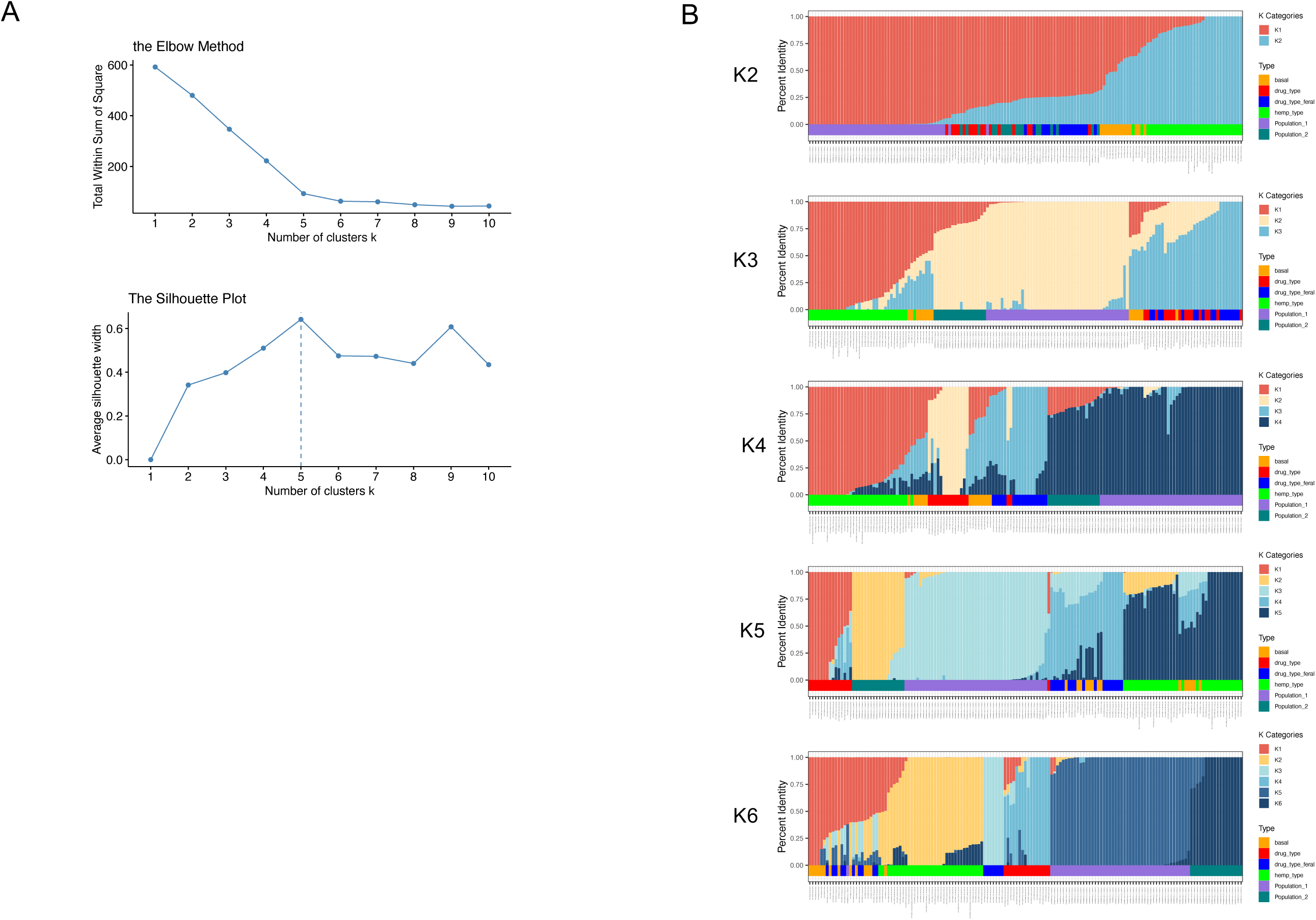
Population structure in the combined datasets **(A)** Optimal K with elbow and silhouette methods **(B)** fastSTRUCTURE analysis for K2-6 using 56,181 SNPs

**Figure S5.**
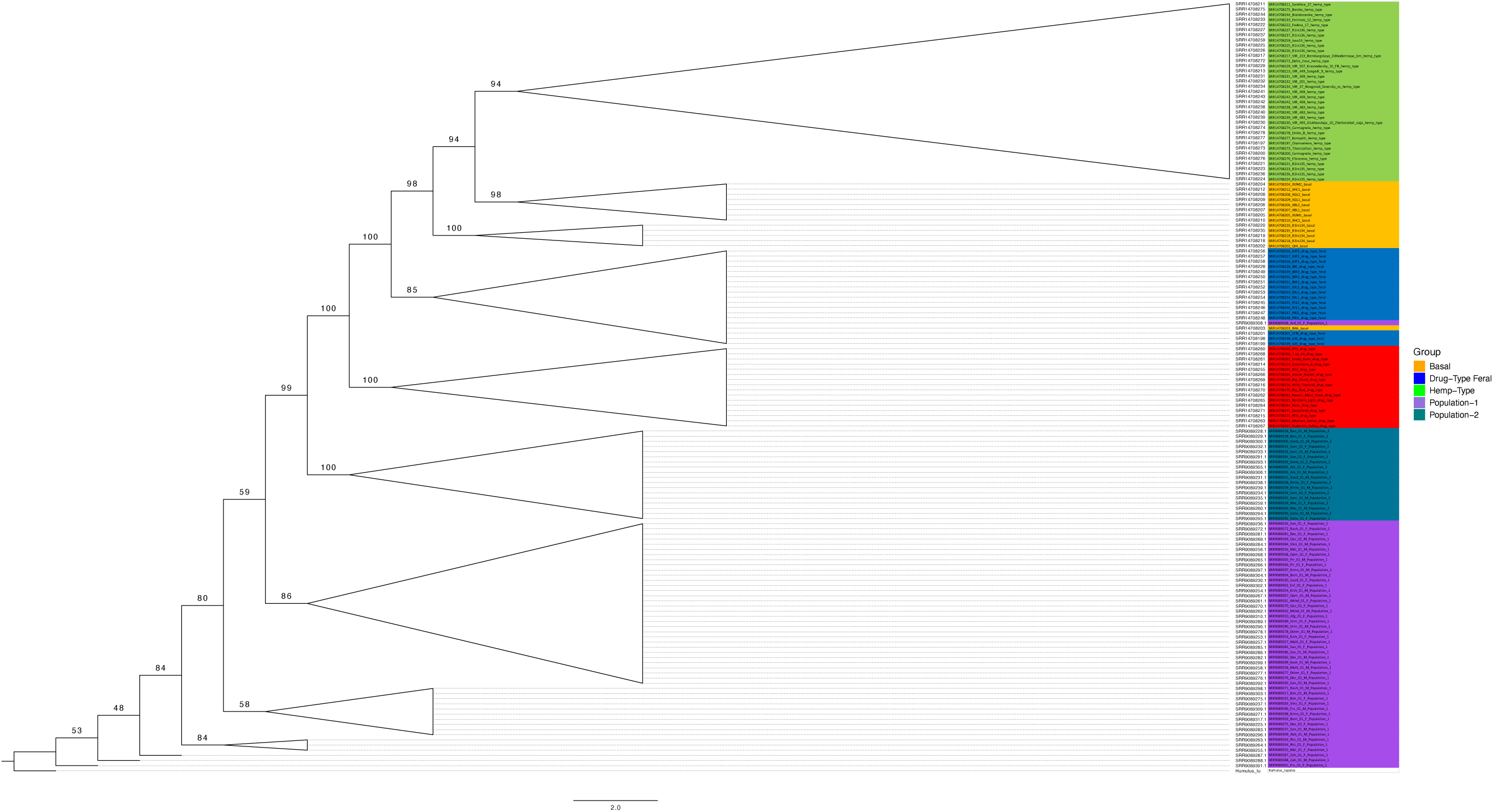
Phylogenetic tree constructed from 13,651 SNPs (no missingness and variant sites only) from the Ren *et al* 2021 and Soorni *et al* 2017 datasets combined rooted using *Humulus lupulus*.

**Figure S6.**
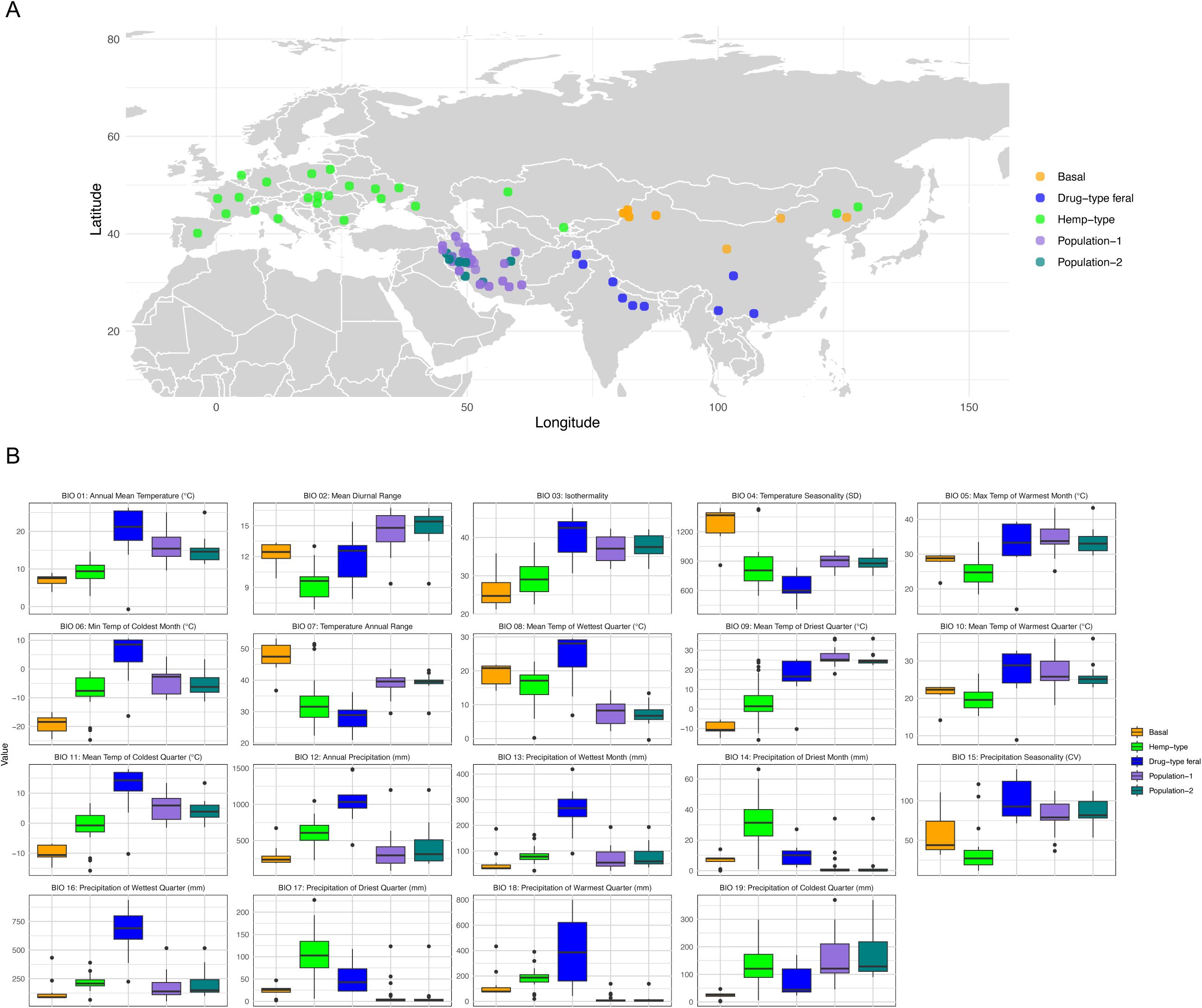
Environmental coverage within the dataset **(A)** Geographic range of 111 samples coloured by population **(B)** Boxplots for the environmental ranges (raw data) for the 111 samples coloured by their phylogenetic association for the 19 bioclimatic variables from WorldClim.

**Figure S7.**
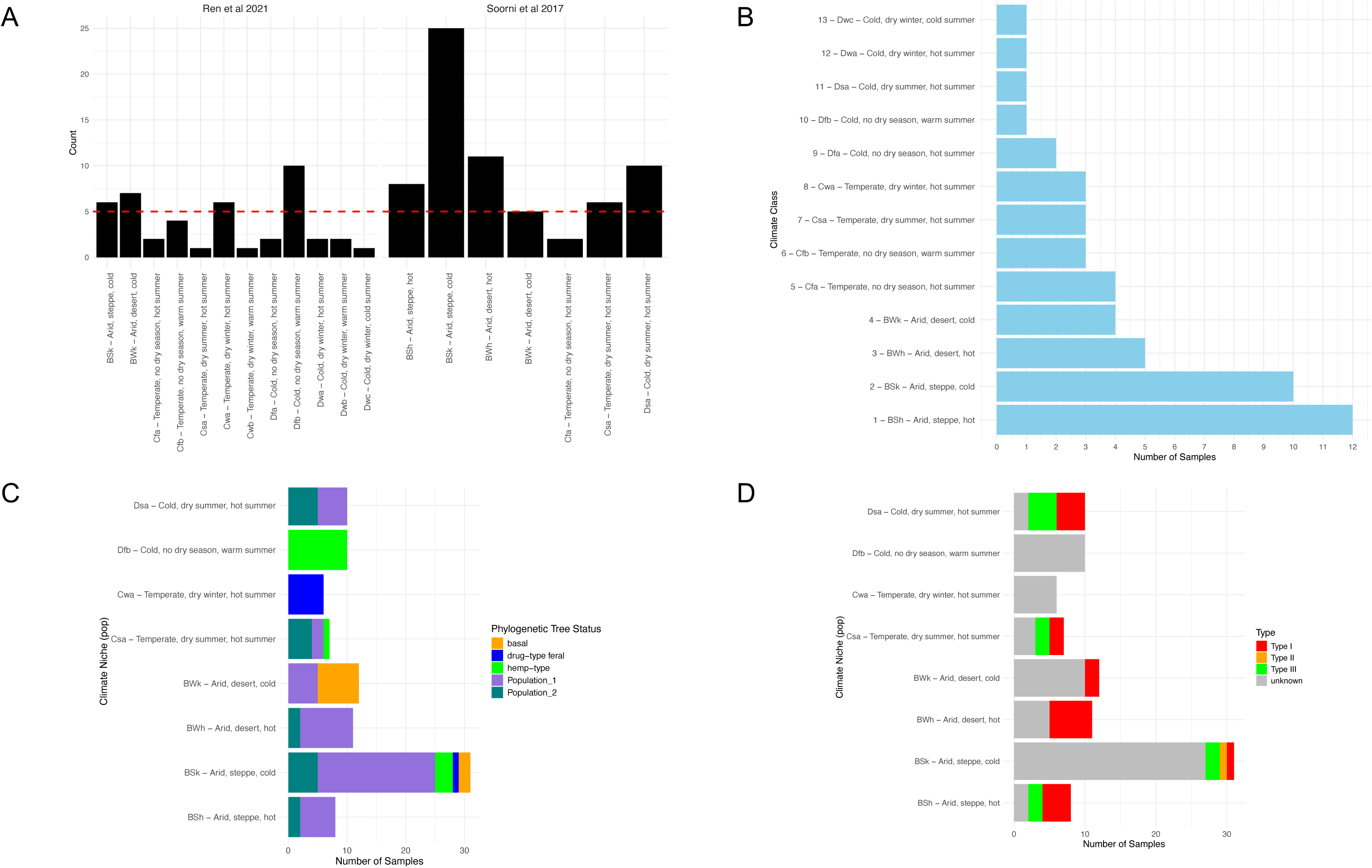
Representation of Köppen-Geiger climate classes by dataset **(A)** Climate classes represented in the combined dataset. Red line indicates climate classes with 5 or greater samples present **(B)** climate classes captured by the core show 13/15 climate niches represented with varying degrees **(C)** Group representation within niches with 5 or greater samples present **(D)** Secondary chemistry where applicable within niches with 5 or greater samples present.

**Figure S8.**
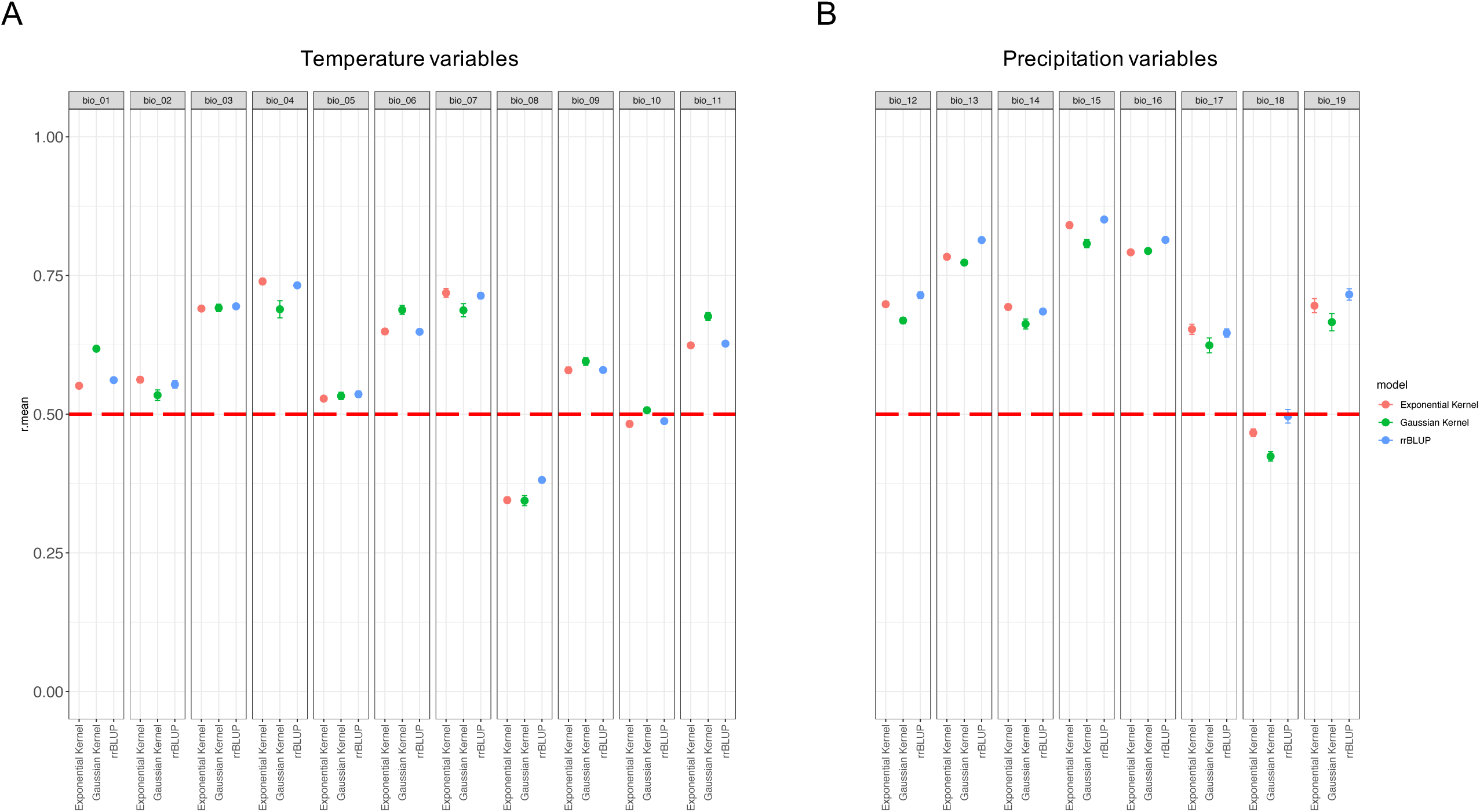
Cross validation showing prediction accuracies for 3 genomic selection models for **(A)** Temperature and **(B)** Precipitation variables from WorldClim for the Ren *et al*., 2021 dataset.

**Figure S9.**
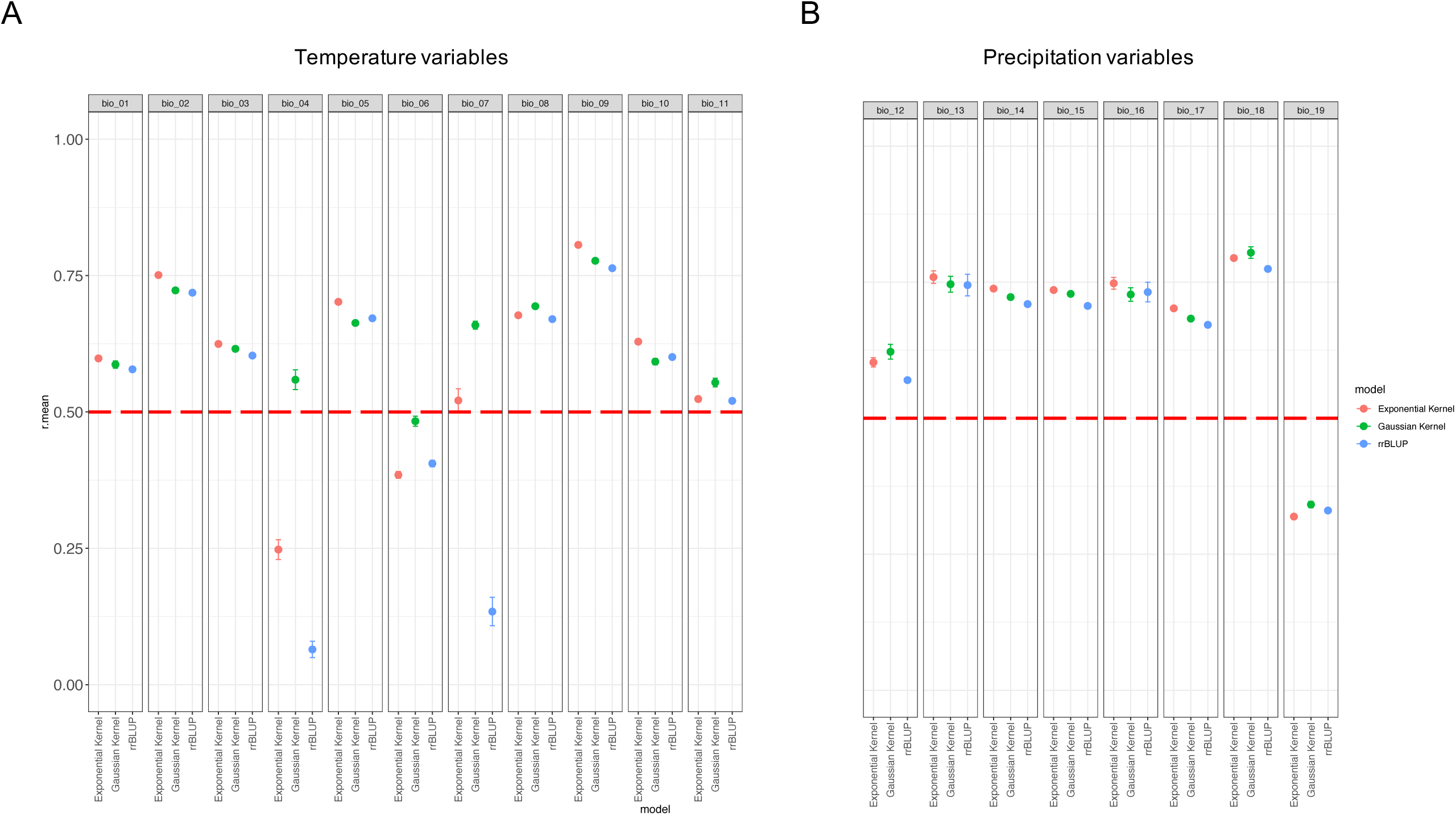
Cross validation showing prediction accuracies for 3 genomic selection models for **(A)** Temperature and **(B)** Precipitation variables from WorldClim for the fusion of the Ren *et al*., 2021 and Soorni *et al*., 2017 datasets.

**Figure S10.**
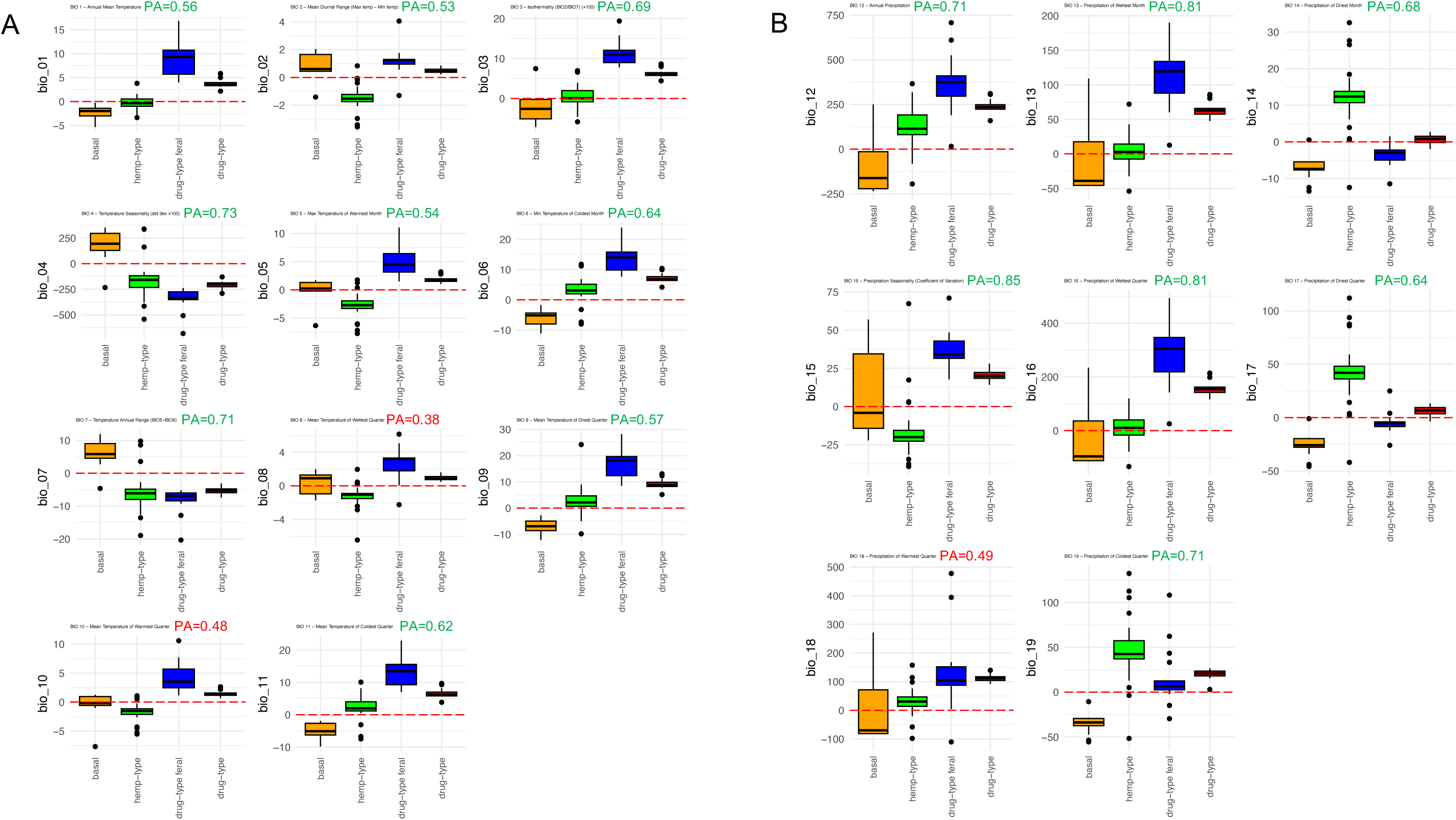
How phylogenetic group in the Ren *et al*., 2021 dataset relates to GEAVs for climate variables **(A)** Temperature **(B)** precipitation. The prediction accuracy (PA) for the rrBLUP model is indicated for each variable.

**Figure S11.**
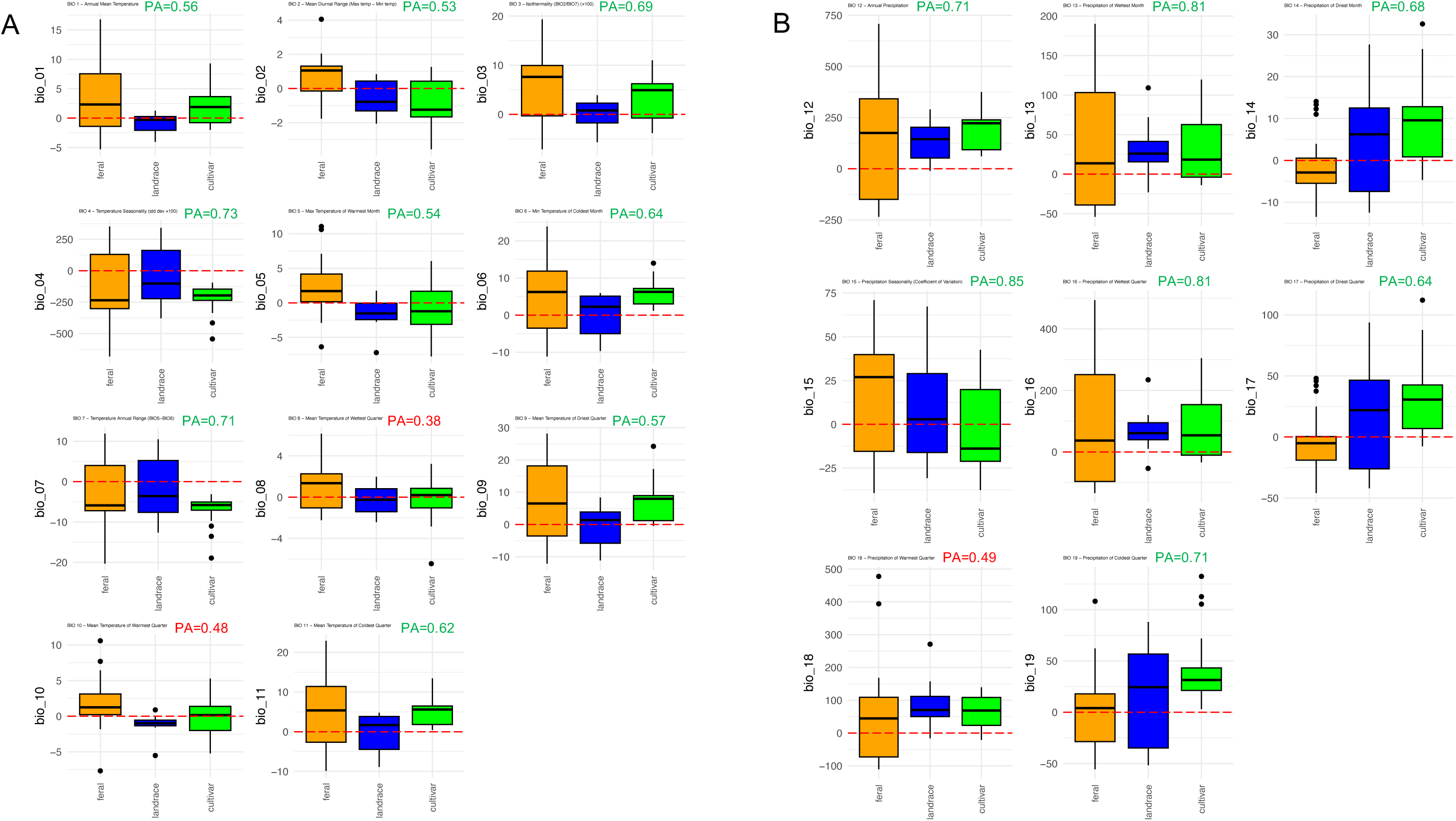
How cultivated status in the Ren *et al*., 2021 dataset relates to GEAVs for climate variables (A) Temperature **(B)** precipitation. The prediction accuracy (PA) for the rrBLUP model is indicated for each variable.

**Figure S12.**
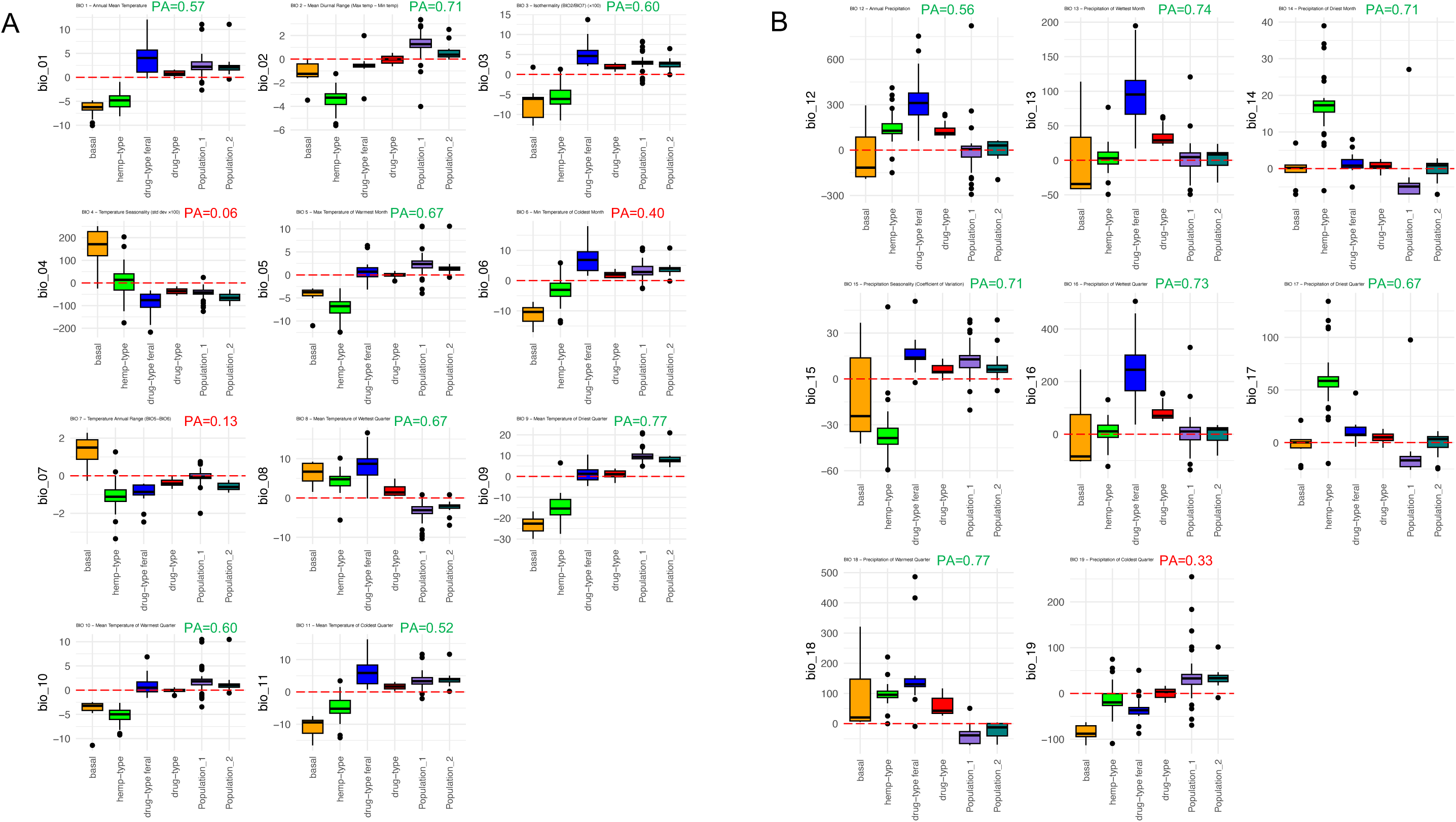
How phylogenetic group relates to GEAVs for climate variables **(A)** Temperature **(B)** Precipitation. The prediction accuracy (PA) for the rrBLUP model is indicated for each variable.

**Figure S13.**
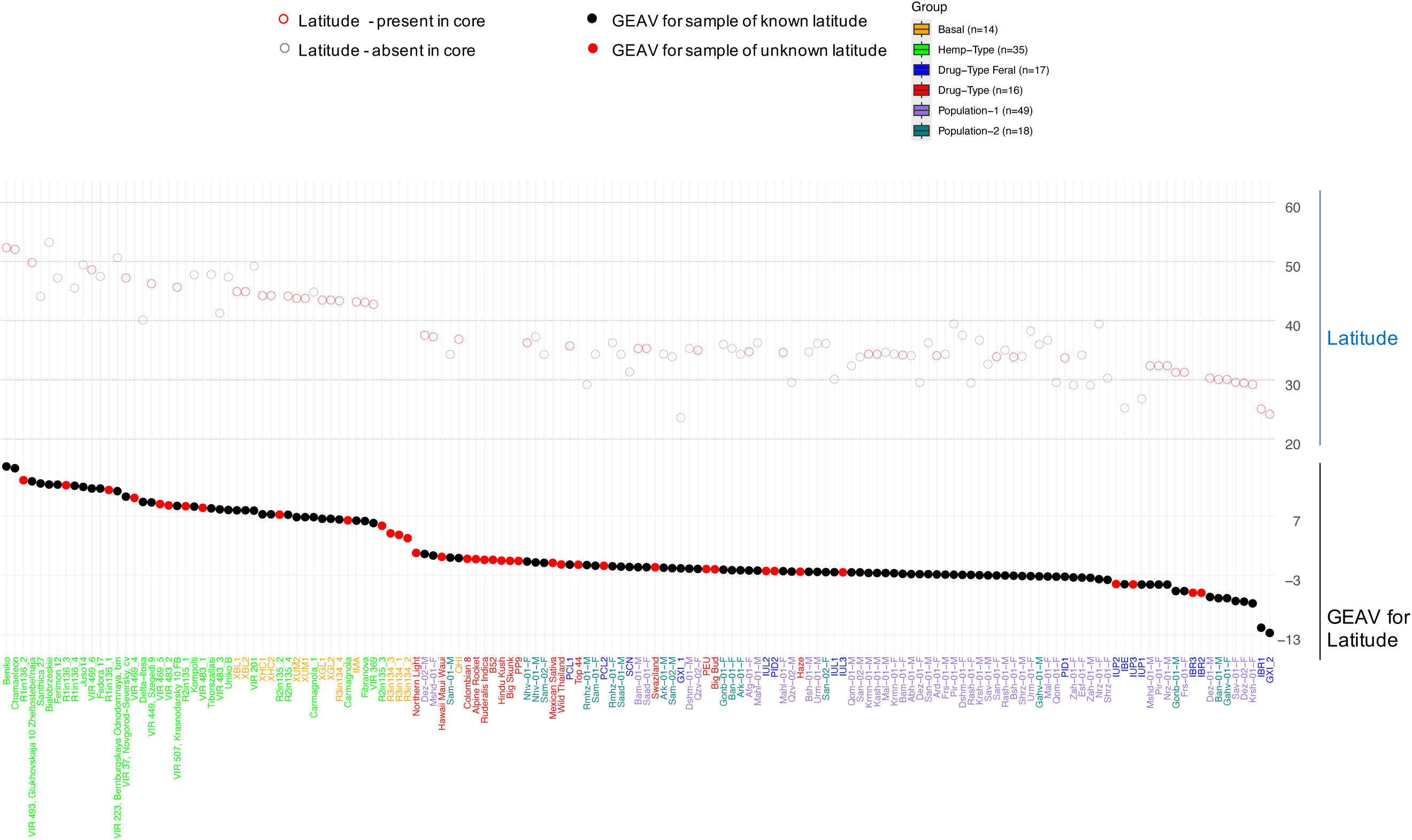
EGS for latitude (PA=0.86) has application in breeding programs for assessing samples of unknown latitudinal origin (red – bottom panel) in particular for the selection of parents with likelihood of matching the latitude of the breeding program which may have implications for flowering time.

**Figure S14.**
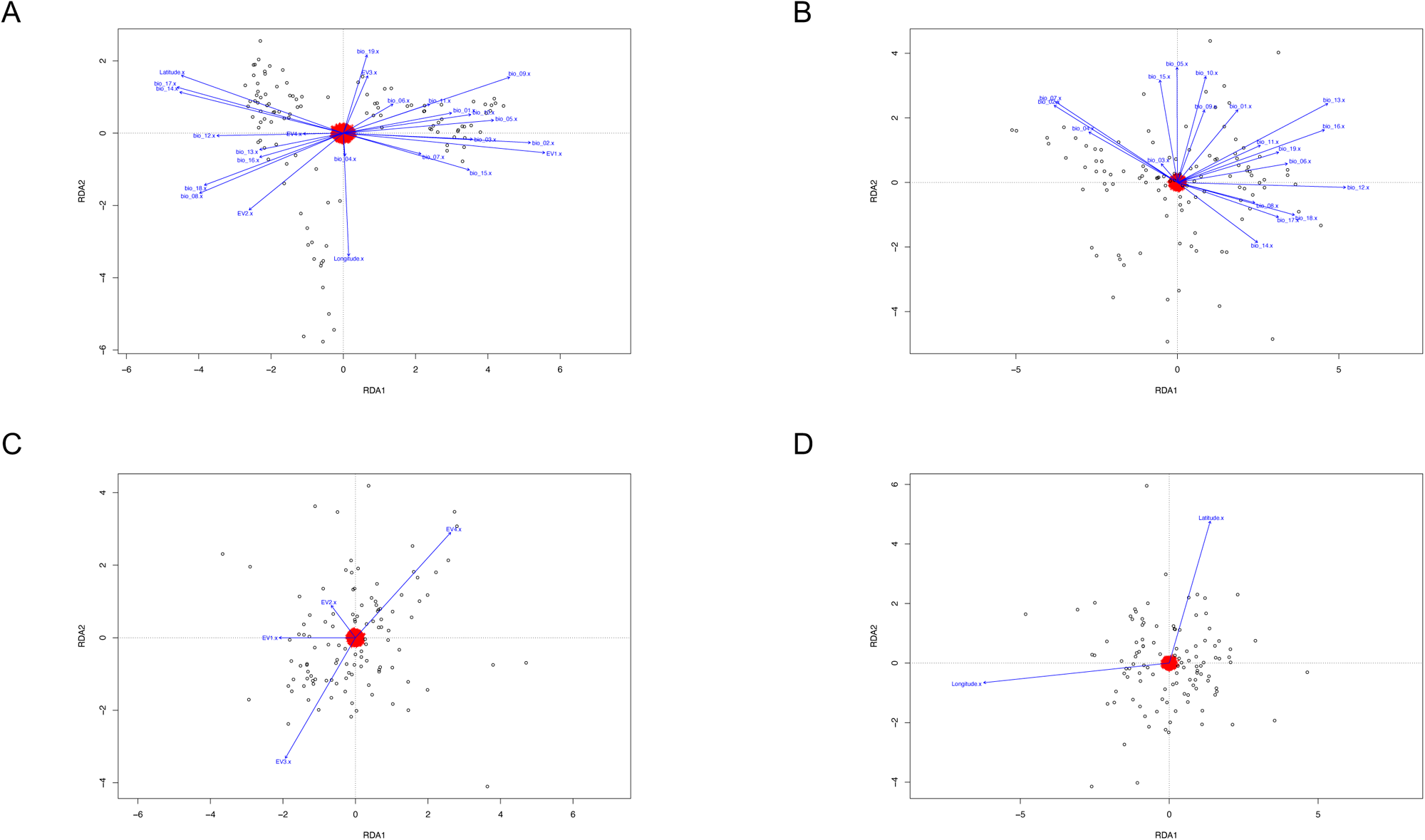
Variation explained by environmental variables for 111/149 samples using Redundancy Analysis (RDA) (**A**) RDA results indicated that 4.3% of genetic variance (adjusted R^2^ = 0.043, p=0.001) was explained by all 19 climate related variables as well as latitude, longitude and population structure (EV1-4) **(B)** Climate alone explained 1.48% of genetic variance (adjusted R^2^ = 0.0148, p=0.001) after controlling for the effects of population structure, latitude and longitude **(C)** Population structure explained 0.03% of genetic variance (adjusted R^2^ = 0.0003, p=0.416) after controlling for the effects of climate, latitude and longitude **(D)** Latitude and Longitude explained 0.35% of genetic variance (adjusted R^2^ = 0.0035, p=0.036) after controlling for the effects of climate and population structure. These and additional RDA results can be seen in Table S8.

**Figure S15.**
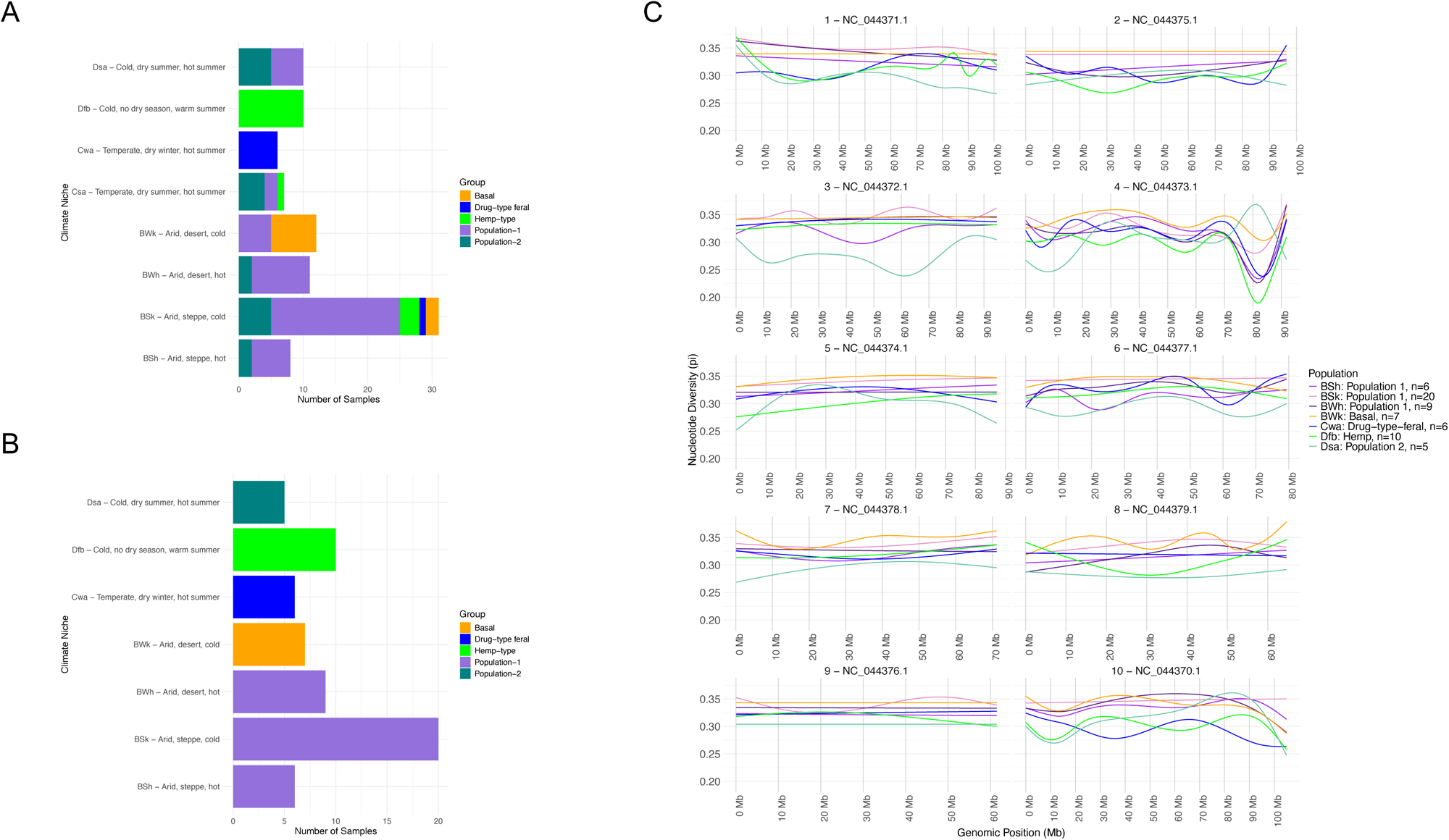
Köppen-Geiger climate classes representation across groups and their relationship with nucleotide diversity **(A)** Group representation within climate classes **(B)** Climate niches which has more then five samples and exclusive to a particular group **(C)** Chromosome level diversity (pi) for seven climate classes using 56,181 SNPs

**Figure S16.**
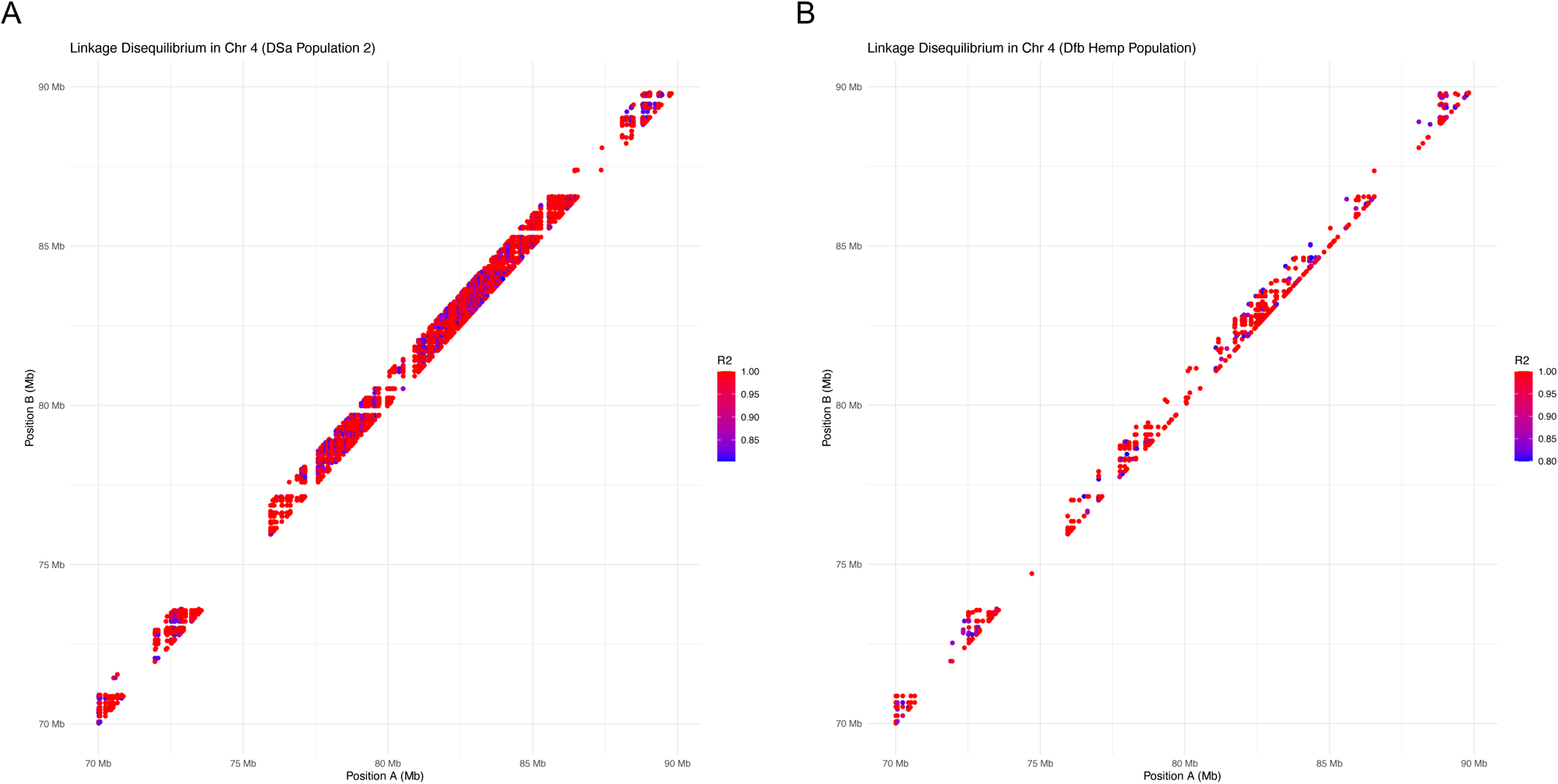
Exploring LD using 1,838 SNPs in the 70Mb to 90Mb region of Chromosome 4 between **(A)** Population-2 in the Dsa climate niche **(B)** Hemp-type in the Dfb climate niche.

**Figure S17.**
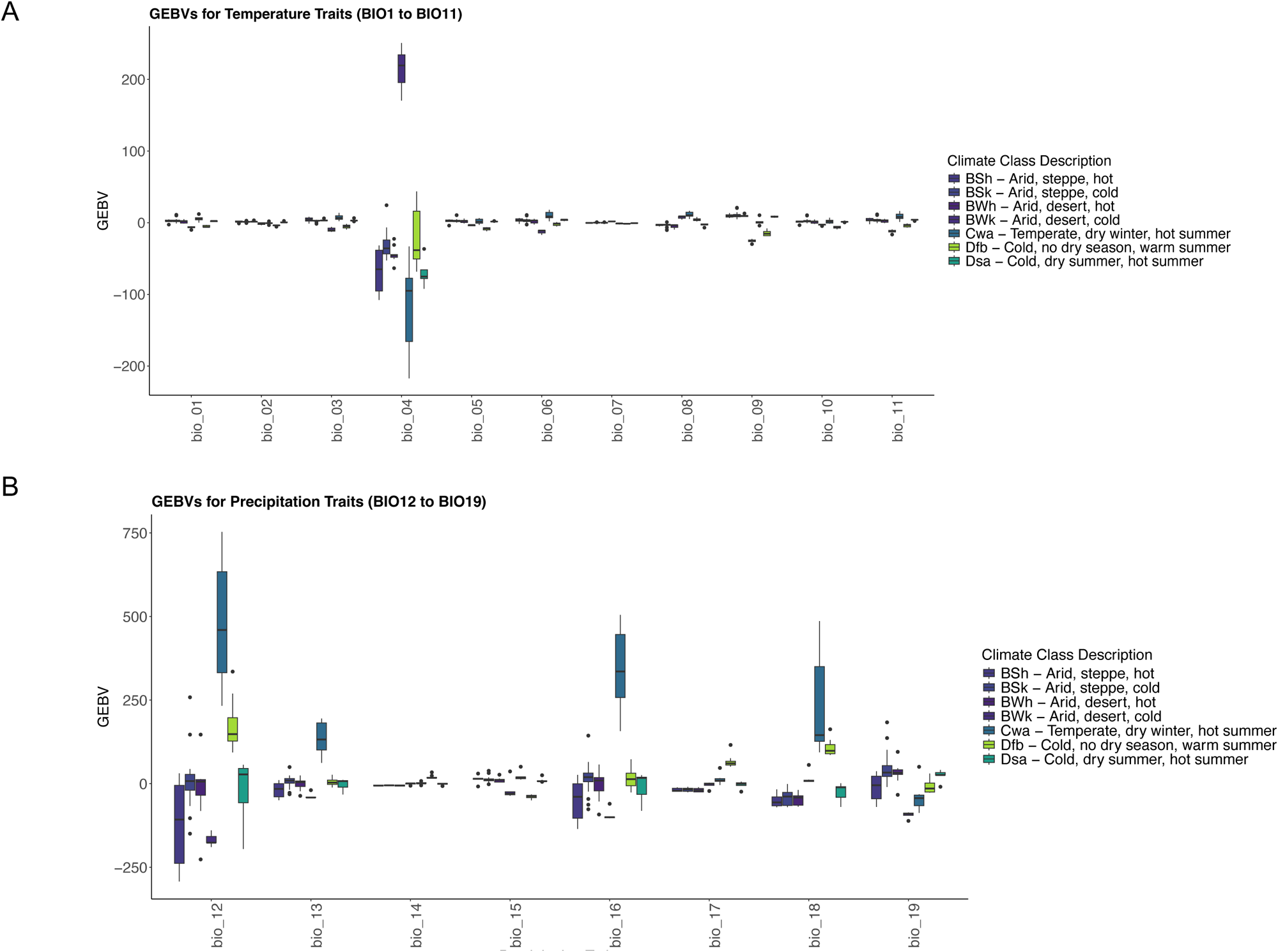
Relationship between climate class and GEAVs for WorldClim climate variables for **(A)** Temperature (BIO1-11) **(B)** Precipitation variables (BIO12-19).

**Figure S18.**
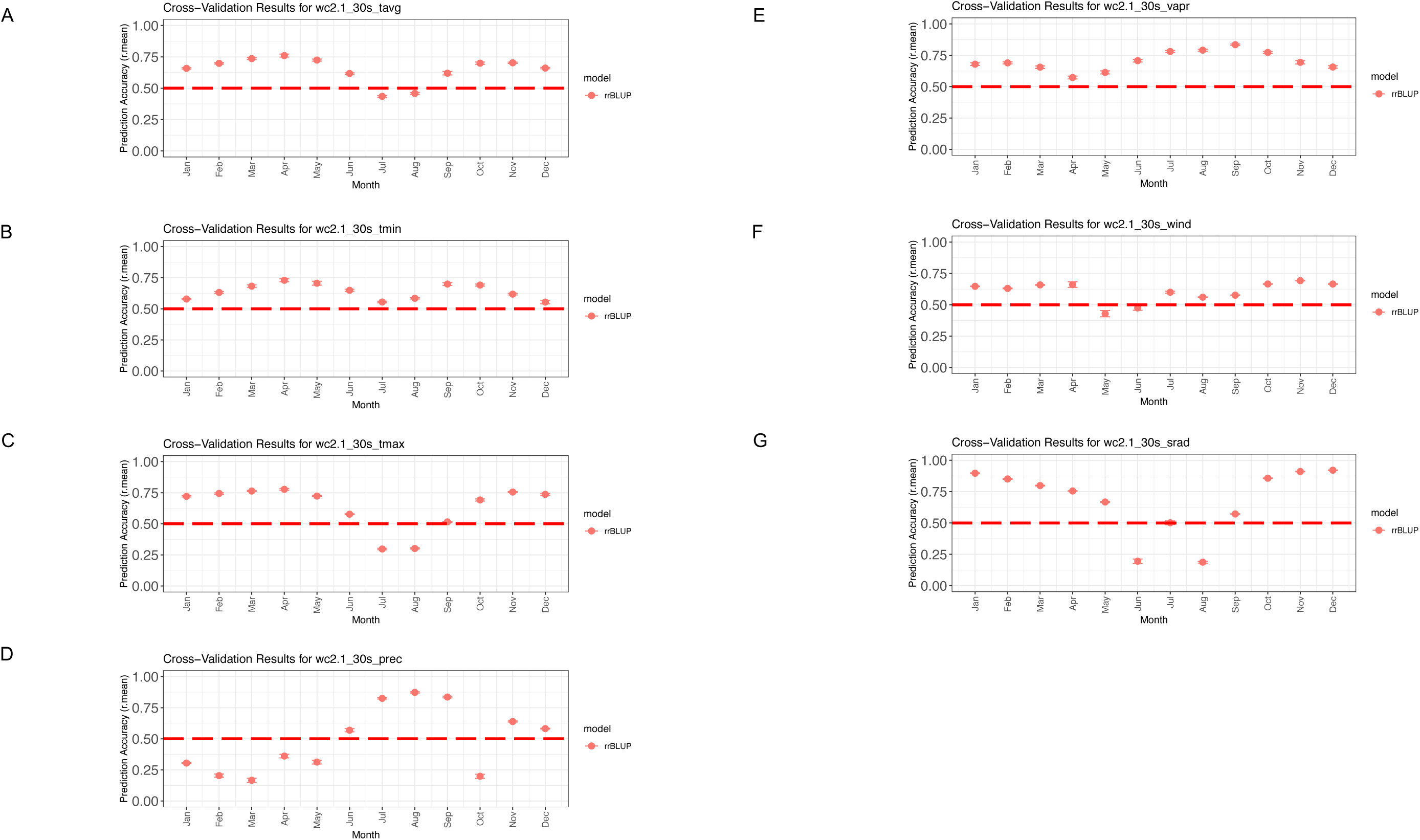
Prediction accuracies for the rrBLUP genomic selection model for the Ren *et al*., 2021 dataset with WorldClim monthly averages (1970-2000) for eight variables **(A)** Average temperature (tavg °C) **(B)** Minimum temperature (tmin °C) **(C)** Maximum temperature (tmax °C) **(D)** Precipitation (mm) **(E)** Water vapor pressure (vapr kPa) **(F)** Wind speed (wind m/s) **(G)** Solar radiation (srad kJ/m^2^/day**) (H)** Elevation (m).

**Figure S19.**
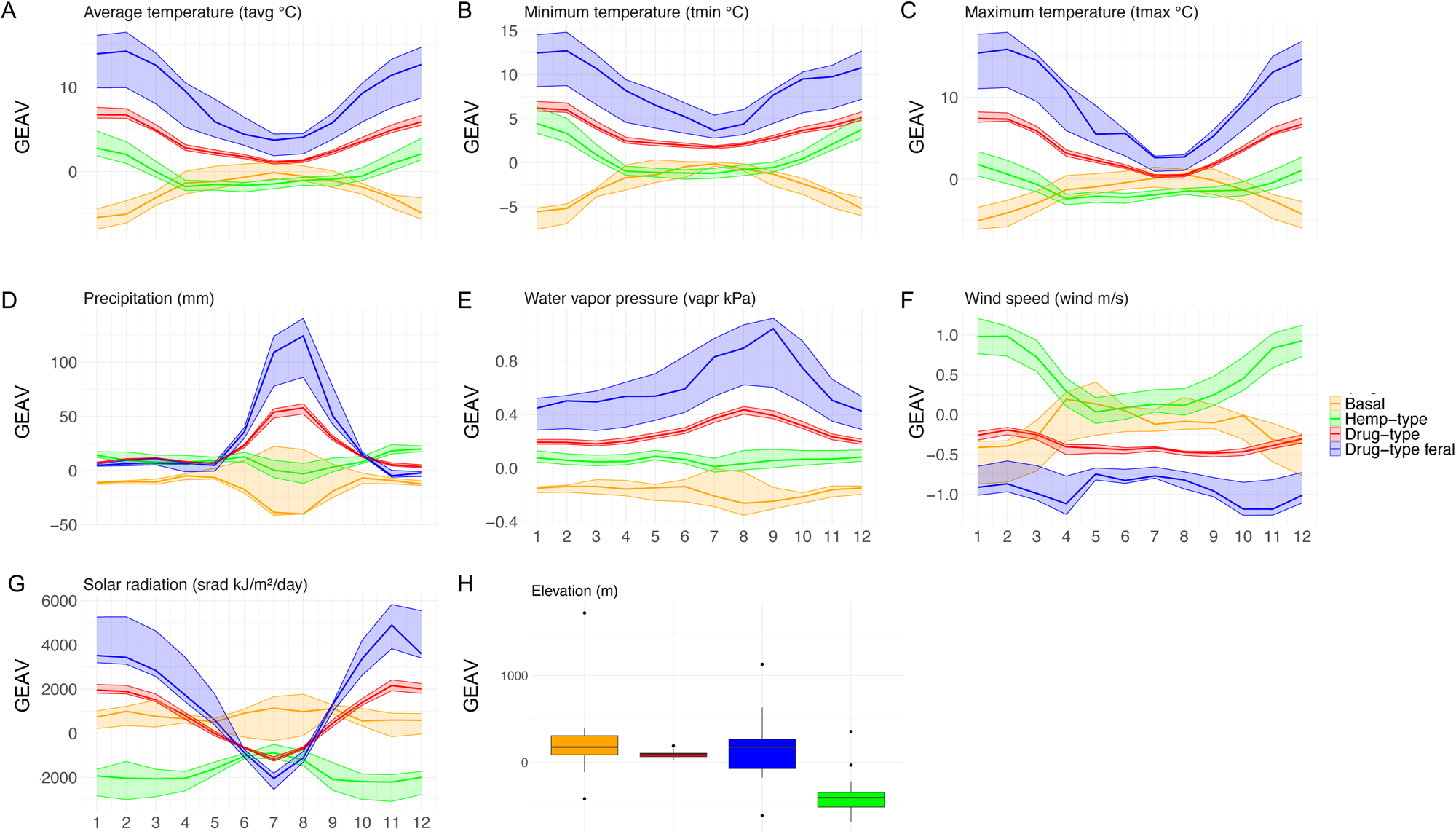
Ren *et al*., 2021 samples coloured by phylogenetic association with the GEAV median, lower quartile (25th percentile) and upper quartile (75th percentile) plotted **(A)** Average temperature (tavg °C) (B) Minimum temperature (tmin °C) **(C)** Maximum temperature (tmax °C) **(D)** Precipitation (mm) **(E)** Water vapor pressure (vapr kPa) **(F)** Wind speed (wind m/s) **(G)** Solar radiation (srad kJ/m^2^/day) **(H)** Elevation (m).

**Figure S20.**
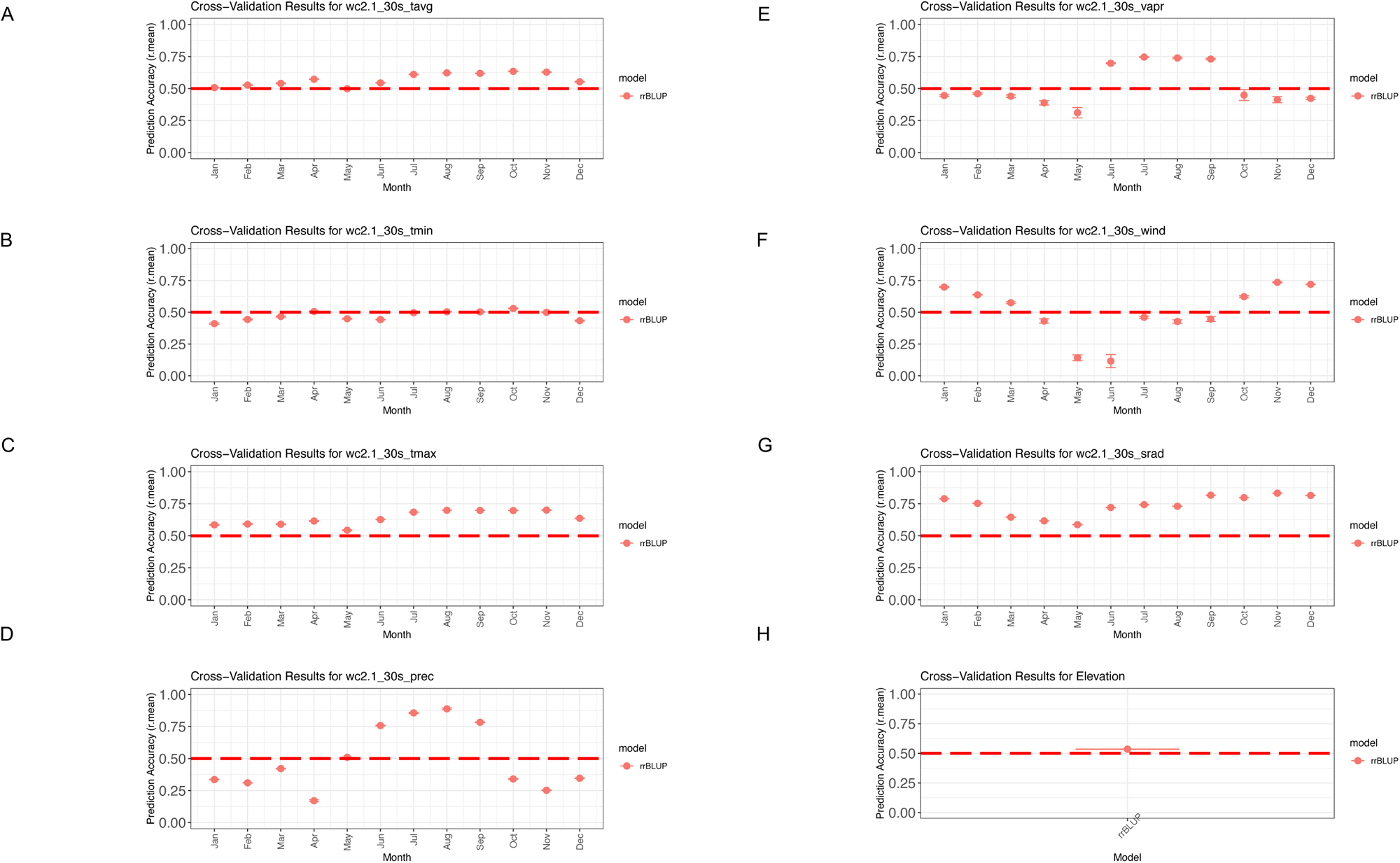
Prediction accuracies for the rrBLUP genomic selection model for the Ren *et al*., 2021 and Soorni *et al*., 2017 datasets with WorldClim monthly averages (1970-2000) for eight variables **(A)** Average temperature (tavg °C**) (B)** Minimum temperature (tmin °C) **(C)** Maximum temperature (tmax °C) **(D)** Precipitation (mm) **(E)** Water vapor pressure (vapr kPa) **(F)** Wind speed (wind m/s) **(G)** Solar radiation (srad kJ/m^2^/day) **(H)** Elevation (m).

**Figure S21.**
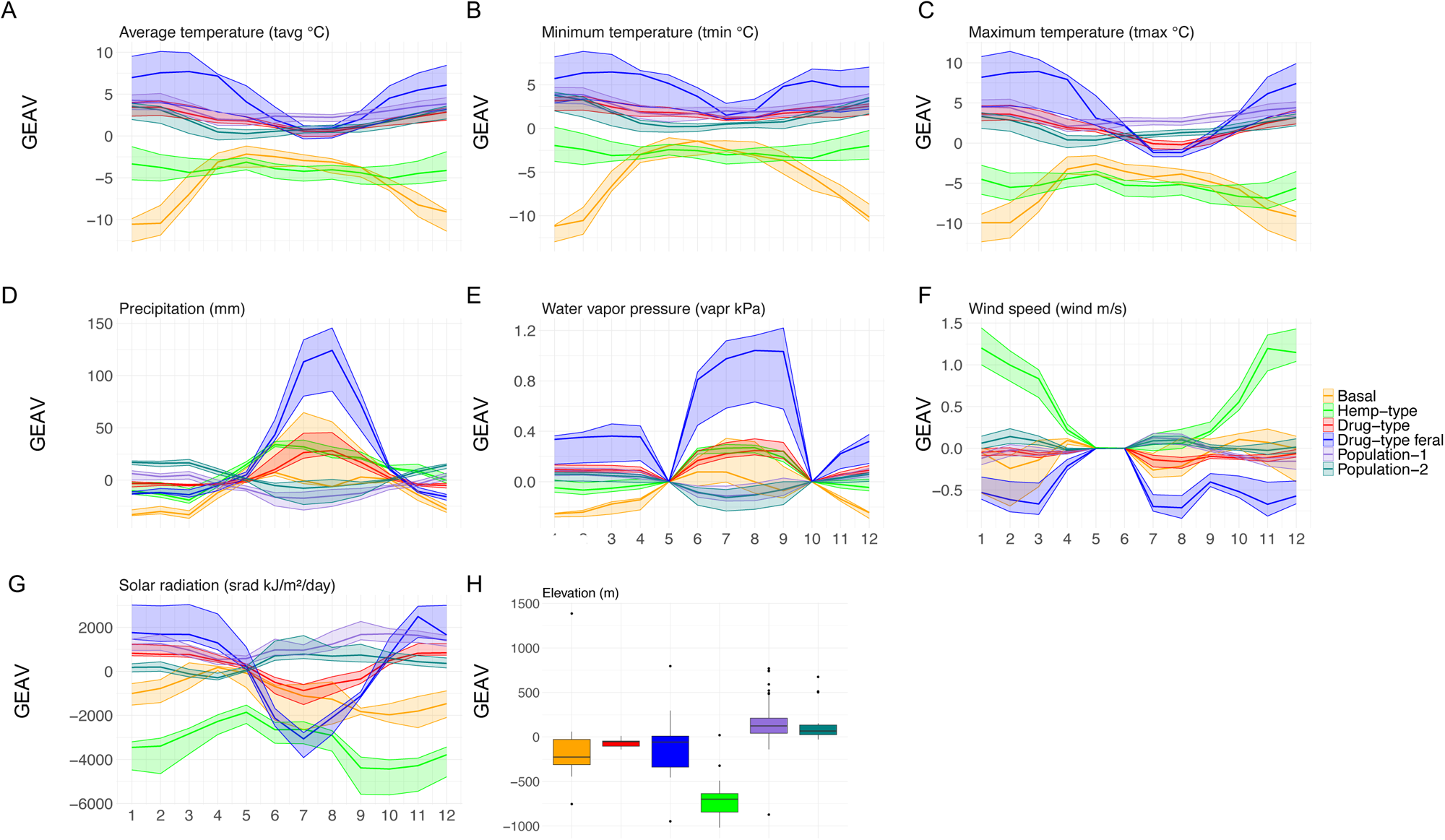
Ren *et al.,* 2021 and Soorni *et al.,* 2017 samples coloured by phylogenetic association with the GEAV median, lower quartile (25th percentile) and upper quartile (75th percentile) plotted for **(A)** Average temperature (tavg °C) **(B)** Minimum temperature (tmin °C) **(C)** Maximum temperature (tmax °C) **(D)** Precipitation (mm) **(E)** Water vapor pressure (vapr kPa) **(F)** Wind speed (wind m/s) (G) Solar radiation (srad kJ/m^2^/day) **(H)** Elevation (m).

**Figure S22.**
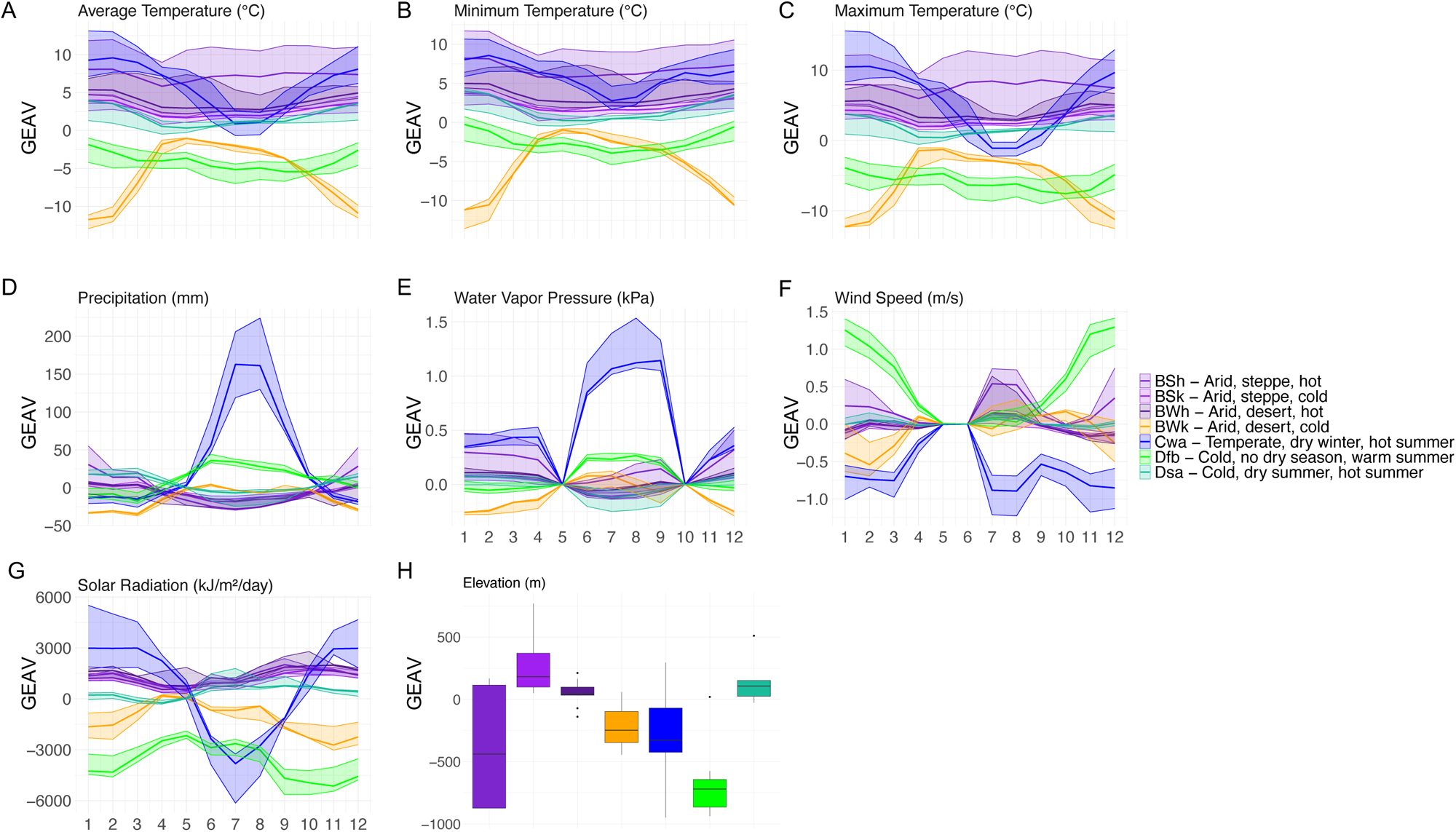
Ren *et al.,* 2021 and Soorni *et al.,* 2017 samples coloured by climate class with the GEAV median, lower quartile (25th percentile) and upper quartile (75th percentile) plotted for **(A)** Average temperature (tavg °C) **(B)** Minimum temperature (tmin °C) **(C)** Maximum temperature (tmax °C) **(D)** Precipitation (mm) **(E)** Water vapor pressure (vapr kPa) **(F)** Wind speed (wind m/s) **(G)** Solar radiation (srad kJ/m^2^/day) **(H)** Elevation (m).

**Figure S23.**
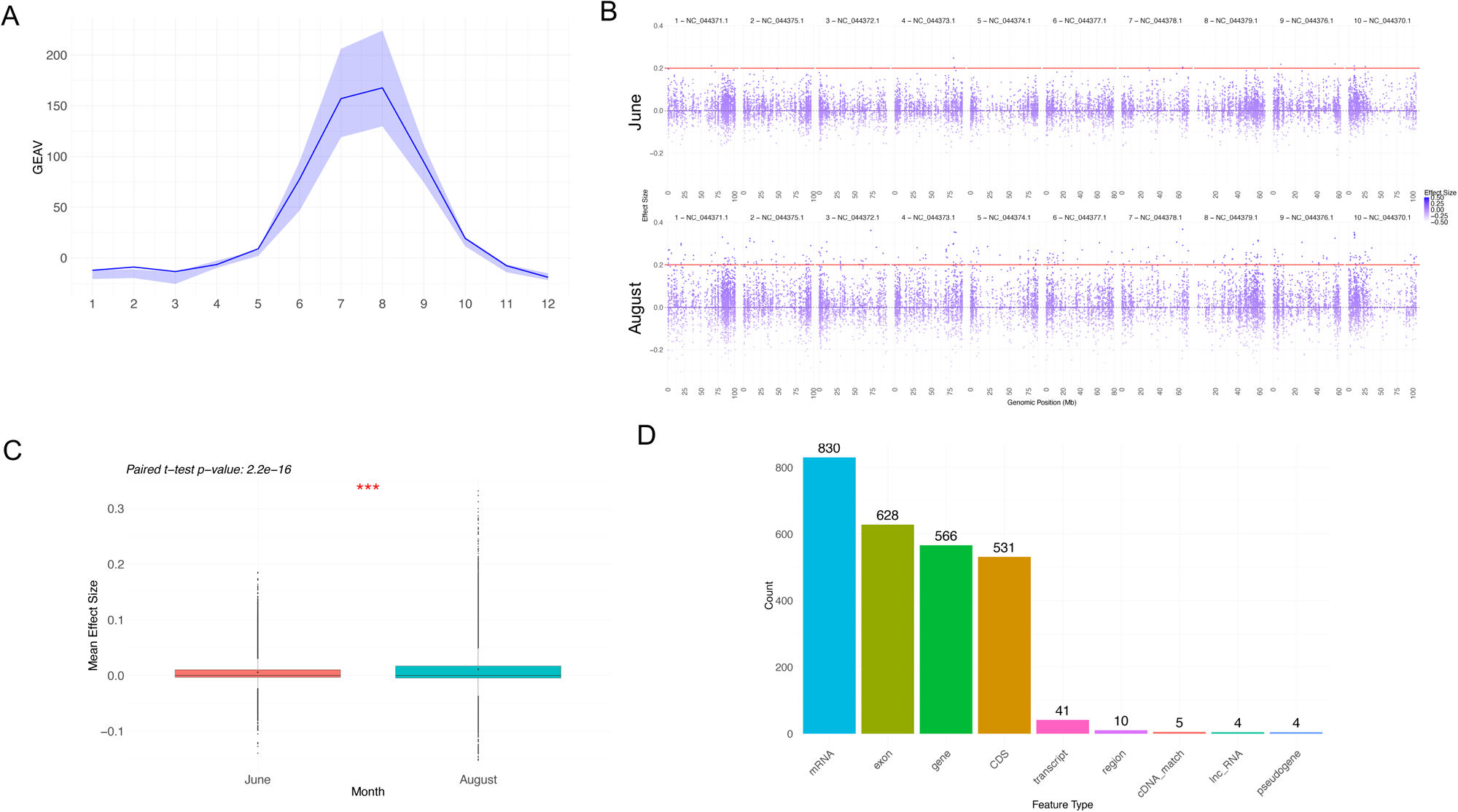
Seasonal response to precipitation in Drug-type feral grouping **(A)** GEAV values for Drug-type feral (n=6) group show seasonal precipitation response. The median line is plotted with the 25^th^ and 75^th^ interquartile range in shaded areas **(B)** Per SNP marker effect size comparison between May and August for six individuals in the Cwa niche **(C)** Paired t-test shows significant difference (p=1.1e-172) between overall mean of maker effect sizes between May and August **(D)** Per SNP paired t-test between January and August results in 788 SNPs with significant change, 566 of which were in gene regions when examining their associations in the cs10 annotation.

**Figure S24.**
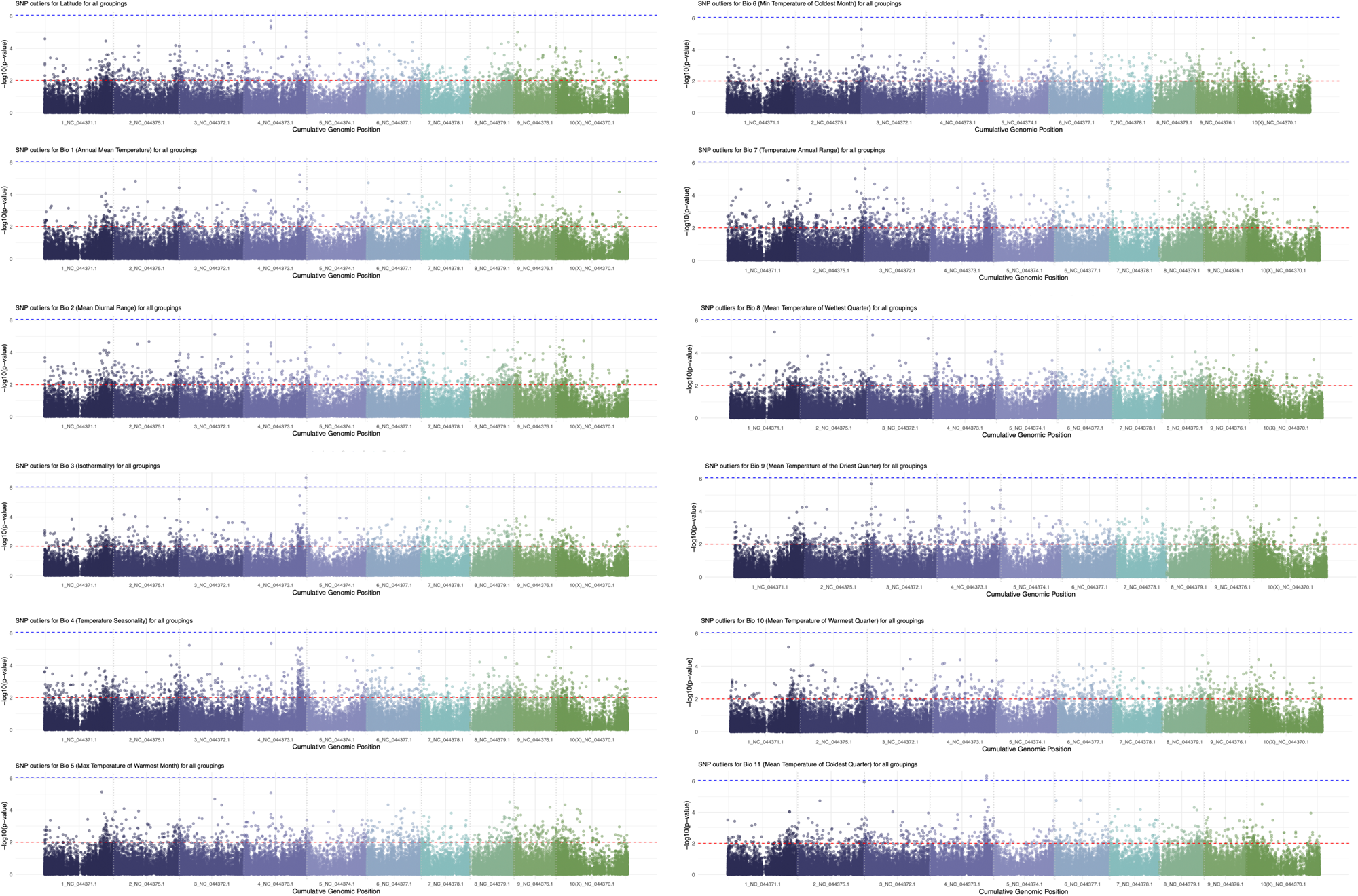
Genome Environment Associations (GEA) for Latitude and WorldClim temperature variables (BIO1-11).

**Figure S25.**
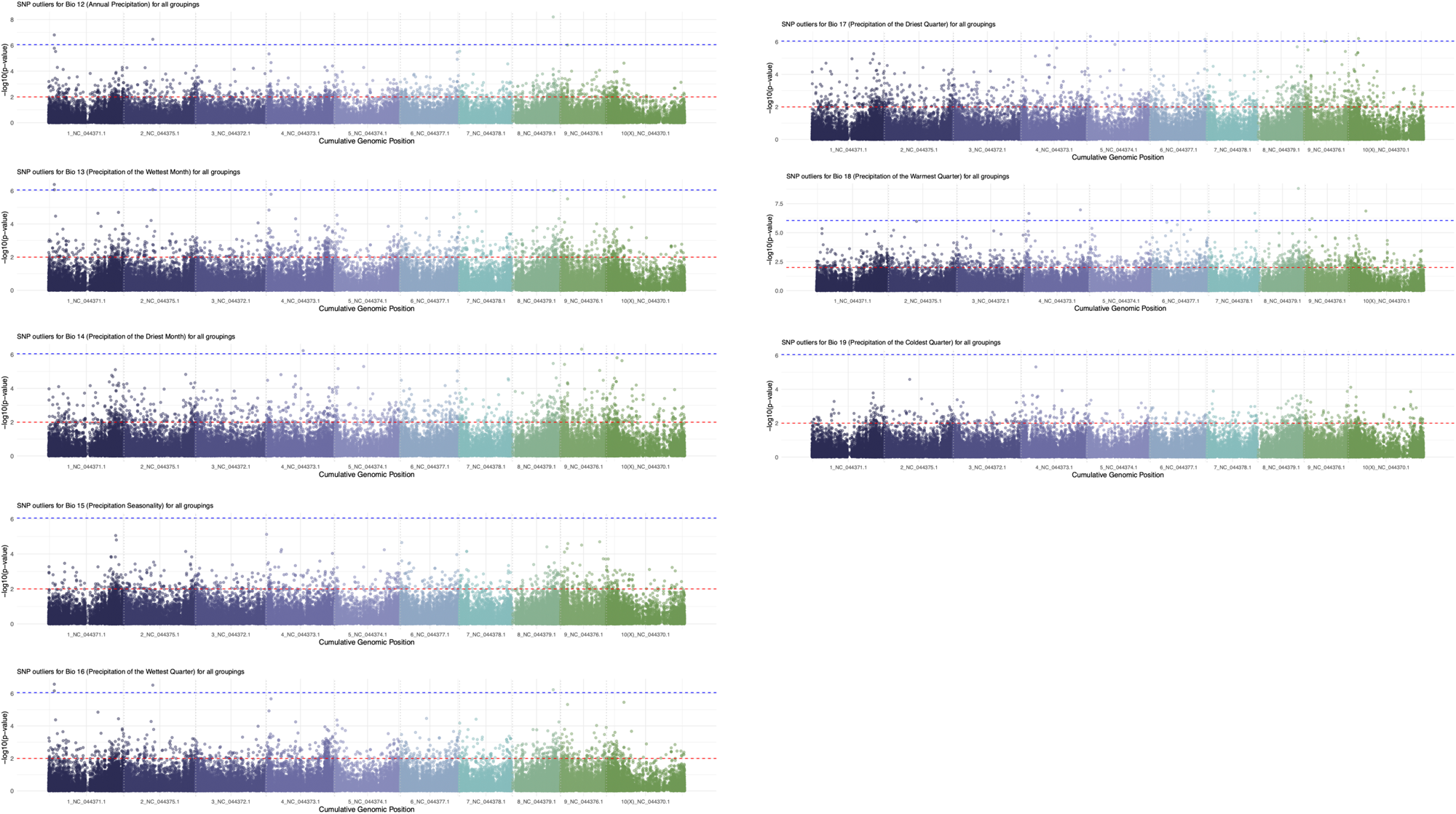
Genome Environment Associations (GEA) for WorldClim precipitation variables (BIO12-19).

**Figure S26.**
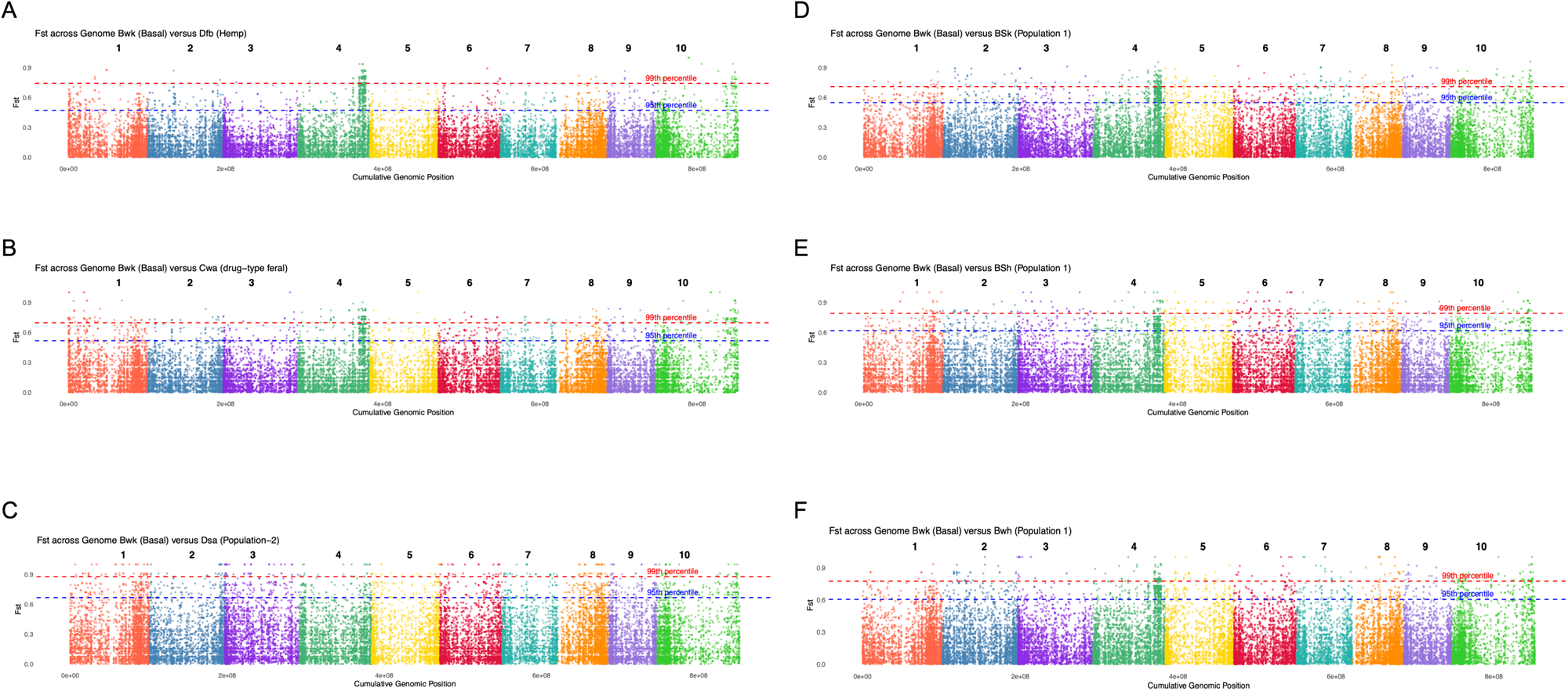
Genome wise Fst comparisons for each population within distinct climate classes against the basal group. Dashed lines indicate the 95^th^ percentile (blue) and the 99^th^ percentile (red). Comparisons are shown between the basal group (in BWk climate class) and population comparisons with **(A)** Hemp-type (in Dfb - Cold, no dry season, warm summer) **(B)** Drug-type feral (in Cwa - Temperate, dry winter, hot summer) **(C)** Population-2 (in Dsa - Cold, dry summer, hot summer) **(D)** Population-1 (in BSk - Arid, steepe, cold) **(E)** Population-1 (in BSh - Arid, steepe, hot) **(F)** Population-1 (in BWh - Arid, desert, hot).

### Table Legends

**Table S1.** Source data from Ren et al., 2021.

**Table S2.** The 44 samples from Ren et al., 2021 which were georeferenced.

**Table S3.** Samples identified as the core (n=25) from the 44 georeferenced samples from Ren et al., 2021.

**Table S4.** GEAV values for each line and the 19 WorldClim bioclimatic traits for 82 samples.

**Table S5.** Source data from Soorni et al., 2017.

**Table S6.** Samples identified as the core (n=50) for the combined Ren and Soorni datasets.

**Table S7.** GEAV values for each line and the 19 WorldClim bioclimatic traits for 149 samples.

**Table S8.** RDA variance partitioning of genetic and structural variation

**Table S9.** Outlier SNPs from GEA analysis (Figure 24/25) and matches in cs10 annotation.

**Table S10.** Results from DAVID analysis.

